# Brain signatures of surprise in EEG and MEG data

**DOI:** 10.1101/2020.01.06.895664

**Authors:** Zahra Mousavi, Mohammad Mahdi Kiani, Hamid Aghajan

## Abstract

The brain is constantly anticipating the future of sensory inputs based on past experiences. When new sensory data is different from predictions shaped by recent trends, neural signals are generated to report this surprise. Existing models for quantifying surprise are based on an ideal observer assumption operating under one of the three definitions of surprise set forth as the Shannon, Bayesian, and Confidence-corrected surprise. In this paper, we analyze both visual and auditory EEG and auditory MEG signals recorded during oddball tasks to examine which temporal components in these signals are sufficient to decode the brain’s surprise based on each of these three definitions. We found that for both recording systems the Shannon surprise is always significantly better decoded than the Bayesian surprise regardless of the sensory modality and the selected temporal features used for decoding.

**Author summary:** A regression model is proposed for decoding the level of the brain’s surprise in response to sensory sequences using selected temporal components of recorded EEG and MEG data. Three surprise quantification definitions (Shannon, Bayesian, and Confidence-corrected surprise) are compared in offering decoding power. Four different regimes for selecting temporal samples of EEG and MEG data are used to evaluate which part of the recorded data may contain signatures that represent the brain’s surprise in terms of offering a high decoding power. We found that both the middle and late components of the EEG response offer strong decoding power for surprise while the early components are significantly weaker in decoding surprise. In the MEG response, we found that the middle components have the highest decoding power while the late components offer moderate decoding powers. When using a single temporal sample for decoding surprise, samples of the middle segment possess the highest decoding power. Shannon surprise is always better decoded than the other definitions of surprise for all the four temporal feature selection regimes. Similar superiority for Shannon surprise is observed for the EEG and MEG data across the entire range of temporal sample regimes used in our analysis.

## Introduction

The predictive coding framework [1] states that the brain is constantly predicting its next sensory input. Past inputs are used by the brain to form prior knowledge in the Bayesian brain model [2, 3]. In fact, the results of brain functions such as perception, eliminating ambiguity, attention, and decision making are dependent on the way the current sensory input and the knowledge gained from previous experiences are combined in the hierarchical inference model of the brain [1] (for review see [4]).

An input different from what the brain has predicted will be surprising in that it generates a form of response measurable by brain imaging techniques. This surprise has been quantified in the literature as a variable representing the error in predicting the future stimuli based on the expectation of the near-optimal observer [5–15]. Studying the characteristics of surprise plays an important role in understanding how learning happens in the brain. In this context, surprise is often represented by a parameter that the brain attempts to minimize during the process of learning [14, 16–18]. A remarkable observation is that the unpredictability of the stimulus, which leads to a high value of surprise, elicits the attention of the observer [6, 7], and produces distinct brain responses [9, 12, 19].

The Shannon surprise [10] has been widely used as a measure for quantifying surprise based on the likelihood of the data [9, 11–13, 15, 19, 20]. Faraji et al. (2018) introduce the notion of “puzzlement” surprise for when the brain encounters an input not matching the model it has created for the world, and consider the Shannon surprise as a quantification of this surprise. They also introduce an alternative quantification of the puzzlement surprise, named the Confidence-corrected surprise, by comparing the posterior distribution of the input with that of a naïve observer (who bases his model on the most recent input and a uniform prior). To the scope of our knowledge, it is not yet examined whether this new definition better corroborates experimental data than the Shannon surprise does.

After the brain has utilized the new input to update its model of the world, an “enlightenment” surprise ensues [14]. The Bayesian surprise, based on comparing the estimated generative distribution of the received stimuli before and after the arrival of each input, was introduced by Baldi (2002) and has been used thereafter by many researchers [9, 12, 18, 67]. The Bayesian surprise can be regarded as a model for describing the “enlightenment” surprise [14].

Bayesian surprise, also called shifts in beliefs or model updating, is believed to map to a completely different region of the human brain from Shannon surprise. Several recent reports based on fMRI (Functional Magnetic Resonance Imaging) data posit that Shannon and Bayesian surprises modulate different brain regions [21–23]. In other words, Shannon and Bayesian surprise definitions model distinct cognitive processes in the brain, with Shannon surprise reflecting the unlikeliness of the input, and Bayesian surprise representing model updating.

In the context of Event-Related Potential (ERP) experiments, temporal components such as the Mismatch Negativity (MMN) and P300 have been widely believed to manifest underlying surprise events in the brain, and hence, reflect the brain’s error in anticipating the next sensory input [24–29]. These surprise-related components have been often described by a stimulus-referenced information-theoretical surprise in experiments in which subjects are presented with a sequence of stimuli randomly selected from a finite set [9, 12, 13]. Abnormal values in these components have also been proposed as biomarkers for some cognitive disorders such as Schizophrenia and Alzheimer’s disease [30–33], reflecting their importance not only in understanding the behavior of the normal brain in handling surprise, but also in the detection of a number of brain disorders. Similar surprise-related temporal components have been reported in the studies of MEG (Magnetoencephalography) signals [34–36].

An experiment specifically designed to account for the different brain reactions to rare and frequent stimuli is the oddball task, in which a randomly composed sequence of standard and deviant stimuli is presented to a subject [27, 28, 37].

Previous surprise modeling studies mainly base their conclusions on a single component extracted from the brain signals, with the MMN [27] or the P300 [12, 24, 25] or both [29] serving as the main such target components. While such single component analysis simplifies the ensuing effort to develop an encoder or a decoder for the brain surprise, it ignores the contribution of other temporal components corresponding to different post-stimulus latencies. Recent studies have developed models using the entire temporal signals, assuming that the entire epoch of the response is modulated by the statistical properties of the input sequence [15, 38].

In this paper, we set out to study which of the three mentioned information-theoretic quantifications of surprise, reflecting distinct concepts of unlikeliness (Shannon) or unexpectedness (Confidence-corrected) of the input, and belief updating (Bayesian), each happening in different brain regions, fits better to empirical EEG data of visual or auditory modalities and auditory MEG data to model the brain’s response in an oddball experiment. The derivation of these three information-theoretic surprise values are based on an assumption that the observer tries to learn the probability transition matrix of stimuli. This question falls within the scope of the Bayesian brain theory by seeking out how the brain creates and updates a probabilistic model for the incoming sequence of stimuli as a precursor for predicting future sensory inputs. The paper formulates its definition of surprise assuming two transition probability values between the standard and deviant stimuli to describe the generative distribution of the stimuli sequence [13].

The study is applied to an experimental oddball dataset with separate runs of visual and auditory tasks recorded by a multi-channel EEG system. Also, we have repeated the study on the recorded MEG of another auditory oddball task. The recorded data is used to train the trial-by-trial decoder of Modirshanechi et al. (2019) as a regressor to measure the surprise of each stimulus. Surprise labels for training the decoder are generated from the sequence of stimuli. Three separate decoder systems are hence trained with labels generated by each of the three definitions of surprise, and the decoding powers of these decoders are compared.

For each definition of surprise and each sensory modality, samples of recorded data in each epoch are selected in four different regimes in order to also examine the relative decoding power attained by using different temporal segments:

- Entire epoch: First, the entire recorded epoch is used as input to train the decoder.
- Samples: Second, the value of a single time instance sampled from every epoch is used as the decoder input. The decoding power of the set of decoders, each trained based on a specific time instance, determines the relative importance of single temporal EEG/MEG samples in estimating the stimuli surprise.
- Intervals: Third, the decoder inputs are selected as aggregates of temporal samples from the start of the epoch to the target time instance. This allows the decoder to utilize the dependency among the temporal EEG/MEG samples. In this regime, the earliest time instance in the epoch for which the decoding power surpasses a fixed confidence level is subsequently detected.
- Segments: Finally, in order to compare the middle and late segments of the EEG/MEG record which contain the well-known points of mismatch negativity (MMN) and P300, respectively, two different decoders are trained with each of these segments as input.

The paper sets forth comparative results for all the mentioned surprise decoders, and elaborates on the relative efficacy of the underlying definitions of surprise as well as the importance of the different time components in decoding the surprise of the stimuli.

## Results

The feature vectors extracted from the EEG/MEG data for each subject are used as input samples to train a surprise decoder. The decoder uses the result of applying a definition of surprise to the input stimuli sequence as the label during the training phase. The decoder we use for this analysis was introduced by Modirshanechi et al. (2019), and we modified its input features as well as the surprise labels to fit our analysis with three surprise definitions and four temporal feature selection regimes as described earlier. We provide a brief description of the decoder’s operation to better motivate our results. More details about the decoder can be found in [15].

The decoder mainly consists of two modules of linear regression. A Lasso linear regression method takes the feature matrix *S_N_* _*×p*_ (extracted from the EEG data according to one of the 4 temporal sample selection regimes) as its input, and the label vector *Y_N_* _× 1_ (calculated from the input stimuli sequence according to one of the 3 three definitions of surprise) as its labels. The Lasso regressor aims to minimize the reconstruction error while observing an added sparsity term, and is used to eliminate the input features which might be irrelevant to reconstructing the surprise, and hence help avoid overfitting. To evaluate the trained model with cross-validation, we used the R-squared measure. Noticing that the decoding power is a function of the integration coefficient *w* (the parameter defining the coefficients of the window of integration; see Materials and Methods), we have reported the maximum decoding power across all *w*s for each regression by employing the best integration window coefficients. After the removal of features with zero coefficients by the Lasso regressor, the remaining features were presumed effective in describing the surprise.

Tables 1-3 and Figs 1-8 summarize our results on both the EEG and MEG data, which are described in detail here.

**Table 1.** Decoding power and chance level for the three definitions of surprise for the temporal regimes of Entire epoch and the three Early, Middle, and Late segments.

|  | EEG Visual Task |  |  | EEG Auditory Task |  |  | MEG Auditory Task |  |  |
| --- | --- | --- | --- | --- | --- | --- | --- | --- | --- |
| Decoding Power | Shannon | Confidence-Corrected | Bayesian | Shannon | Confidence-Corrected | Bayesian | Shannon | Confidence - Corrected | Bayesian |
| Entire epoch | 0.482 | 0.331 | 0.264 | 0.375 | 0.278 | 0.198 | 0.134 | 0.070 | 0.033 |
| Early | 0.017 | 0.006 | 0.002 | 0.026 | 0.027 | 0.005 | 0.027 | 0.009 | 0.009 |
| Middle | 0.348 | 0.233 | 0.195 | 0.306 | 0.229 | 0.167 | 0.141 | 0.095 | 0.046 |
| Late | 0.414 | 0.287 | 0.227 | 0.340 | 0.248 | 0.185 | 0.088 | 0.056 | 0.031 |
| Chance Level |  |  |  |  |  |  |  |  |  |
| Entire epoch | -0.0038 ± 0.0046 | -0.0038 ± 0.0049 | -0.0038 ± 0.0050 | -0.0030 ± 0.0044 | -0.0033 ± 0.0043 | -0.0033 ± 0.0043 | -0.0031 ± 0.0050 | -0.0031 ± 0.0051 | -0.0031 ± 0.0050 |
| Early | -0.0038 ± 0.0047 | -0.0038 ± 0.0049 | -0.0035 ± 0.0048 | -0.0031 ± 0.0042 | -0.0033 ± 0.0045 | -0.0032 ± 0.0042 | -0.0031 ± 0.0050 | -0.0031 ± 0.0051 | -0.0031 ± 0.0051 |
| Middle | -0.0038 ± 0.0046 | -0.0038 ± 0.0047 | -0.0038 ± 0.0047 | -0.0028 ± 0.0063 | -0.0033 ± 0.0042 | -0.0031 ± 0.0043 | -0.0031 ± 0.0050 | -0.0031 ± 0.0051 | -0.0031 ± 0.0051 |
| Late | -0.0038 ± 0.0046 | -0.0038 ± 0.0049 | -0.0038 ± 0.0047 | -0.0030 ± 0.0041 | -0.0033 ± 0.0043 | -0.0032 ± 0.0042 | -0.0031 ± 0.0050 | -0.0031 ± 0.0051 | -0.0004 ± 0.0069 |

**Table 2.** P-values of the t-test for comparing three temporal segments of Early, Middle, and Late.

| *EEG Visual Task* | Middle vs. Late | Early vs. Middle | Early vs. Late |
| --- | --- | --- | --- |
| *Bayesian* | 0.33 | 8.77E-10 | 9.32E-14 |
| *Shannon* | 0.17 | 1.89E-11 | 1.05E-16 |
| *Confidence- Corrected* | 0.17 | 1.46E-09 | 1.79E-14 |
| *EEG Auditory Task* |  |  |  |
| *Bayesian* | 0.55 | 5.23E-14 | 4.83E-08 |
| *Shannon* | 0.41 | 1.24E-13 | 8.11E-11 |
| *Confidence- Corrected* | 0.55 | 1.01E-10 | 6.59E-08 |
| **MEG Auditory Task** |  |  |  |
| *Bayesian* | 0.44 | 0.005 | 0.16 |
| *Shannon* | 0.17 | 0.0006 | 0.02 |
| *Confidence- Corrected* | 0.16 | 0.0006 | 0.006 |

Highlights indicate p-values < 0.05.

**Table 3.**
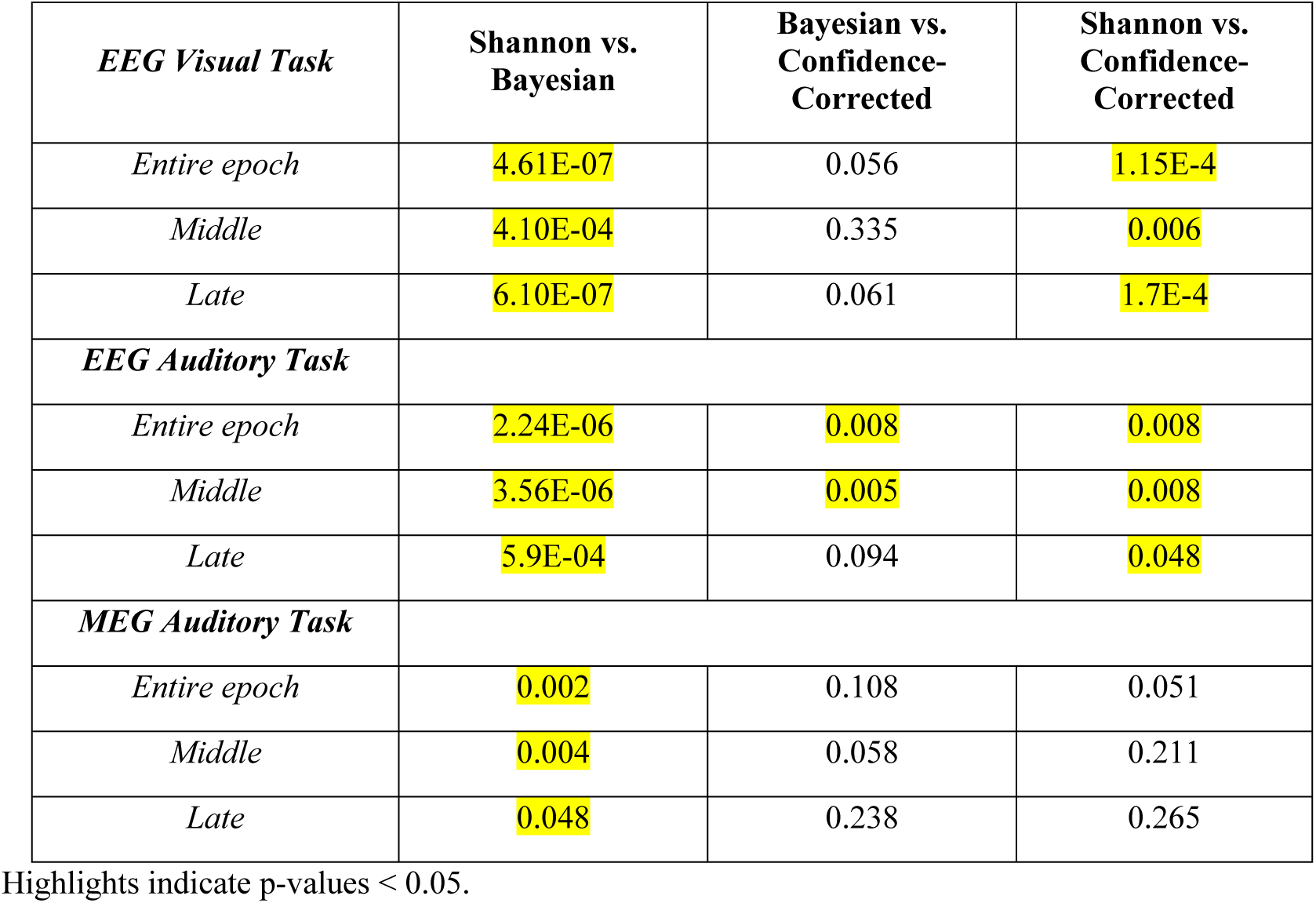
P-values of the t-test of comparing three surprise quantifications.

| **EEG Visual Task** | **Shannon vs. Bayesian** | **Bayesian vs. Confidence-Corrected** | **Shannon vs. Confidence-Corrected** |
| --- | --- | --- | --- |
| *Entire epoch* | 4.61E-07 | 0.056 | 1.15E-4 |
| *Middle* | 4.10E-04 | 0.335 | 0.006 |
| *Late* | 6.10E-07 | 0.061 | 1.7E-4 |
| **EEG Auditory Task** |  |  |  |
| *Entire epoch* | 2.24E-06 | 0.008 | 0.008 |
| *Middle* | 3.56E-06 | 0.005 | 0.008 |
| *Late* | 5.9E-04 | 0.094 | 0.048 |
| **MEG Auditory Task** |  |  |  |
| *Entire epoch* | 0.002 | 0.108 | 0.051 |
| *Middle* | 0.004 | 0.058 | 0.211 |
| *Late* | 0.048 | 0.238 | 0.265 |

Highlights indicate p-values < 0.05.

**Fig 1.**
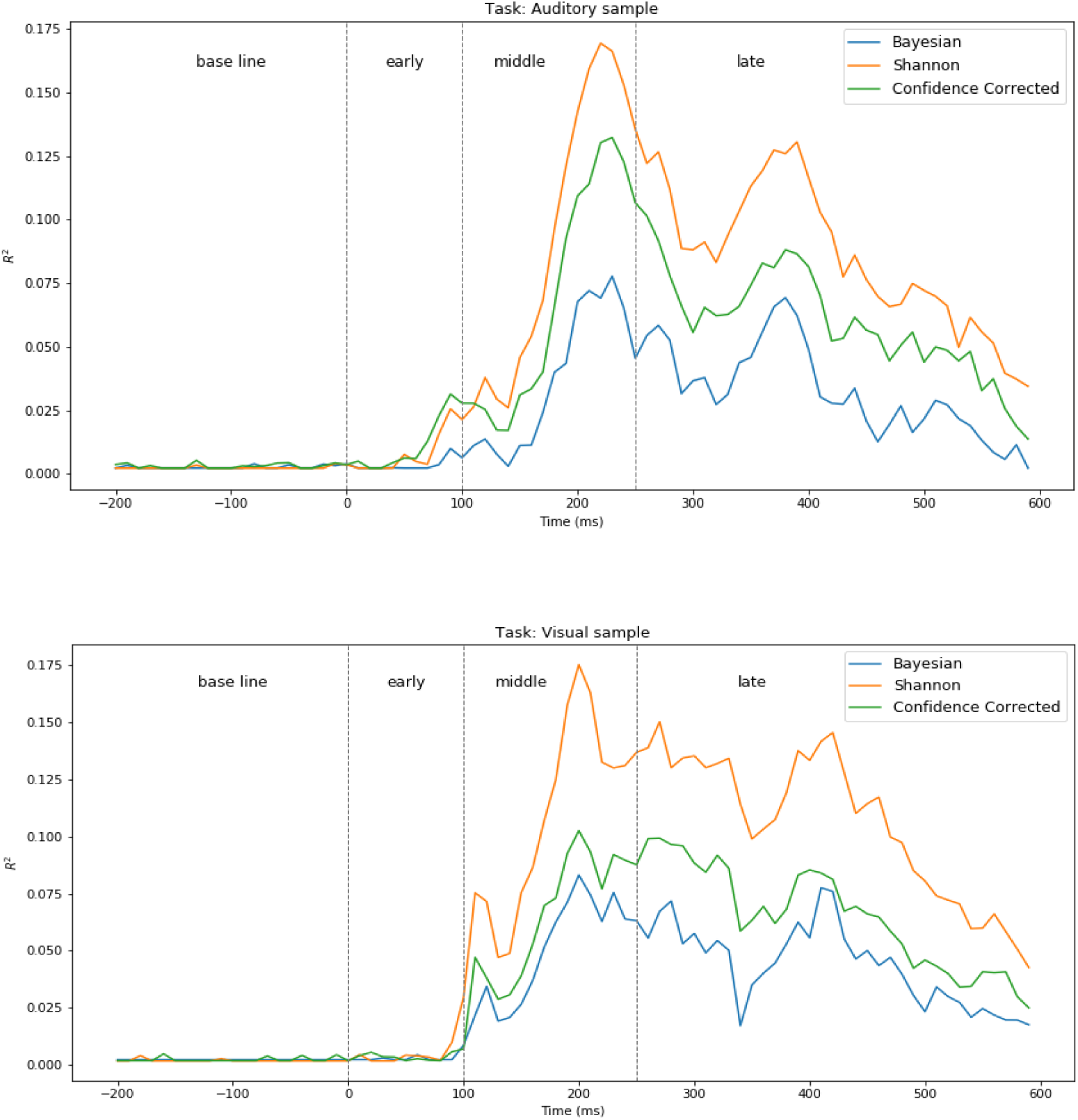
Surprise decoding powers of the three surprise quantifications for every single temporal *Sample* in EEG auditory and visual tasks. Dashed lines indicate selected temporal segments (colors should be used in printing).

**Fig 2.**
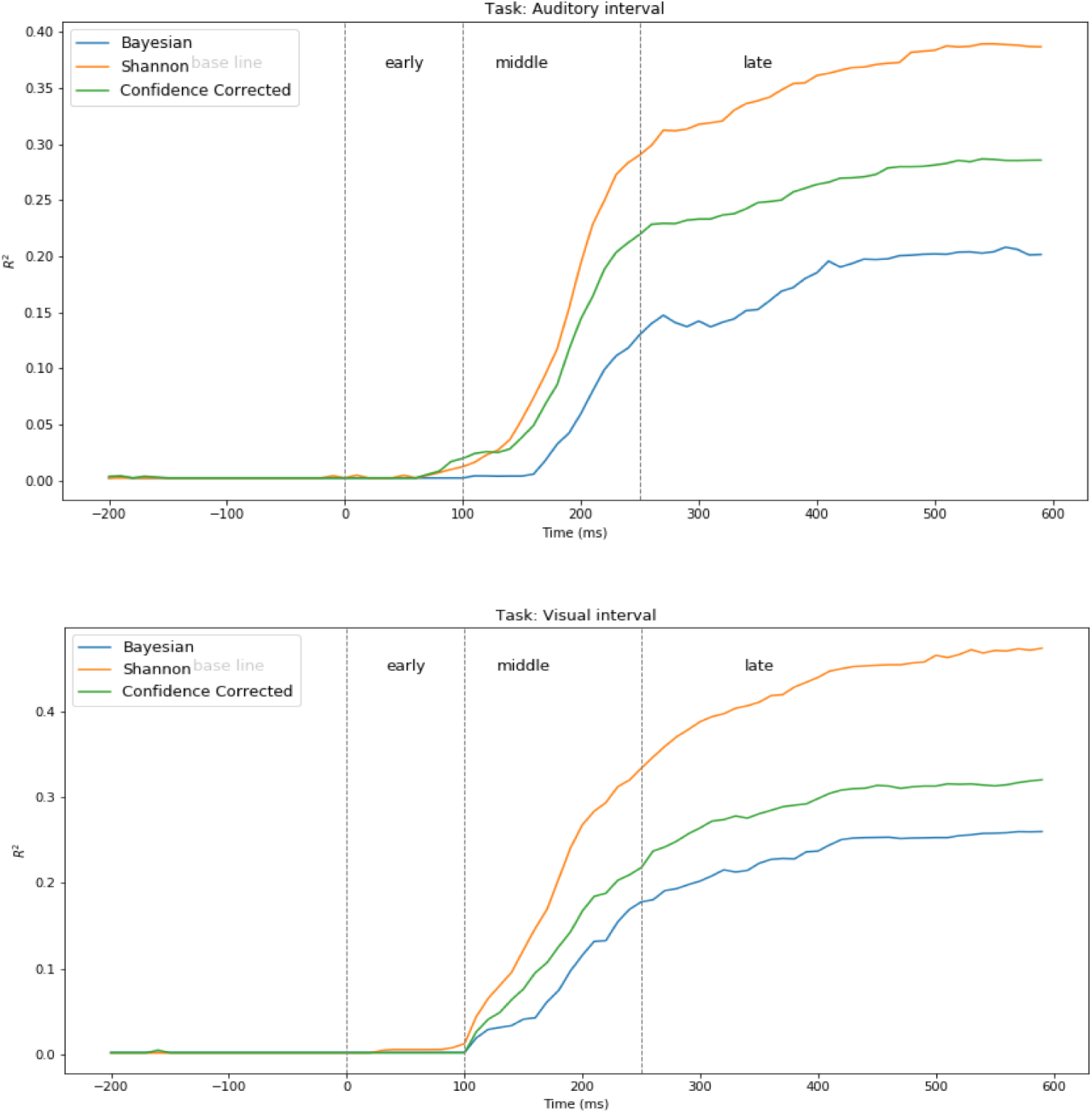
Surprise decoding powers of the three surprise quantifications for the temporal *Interval* [-200ms, t] in EEG auditory and visual tasks. Dashed lines indicate selected temporal segments (colors should be used in printing).

**Fig 3.**
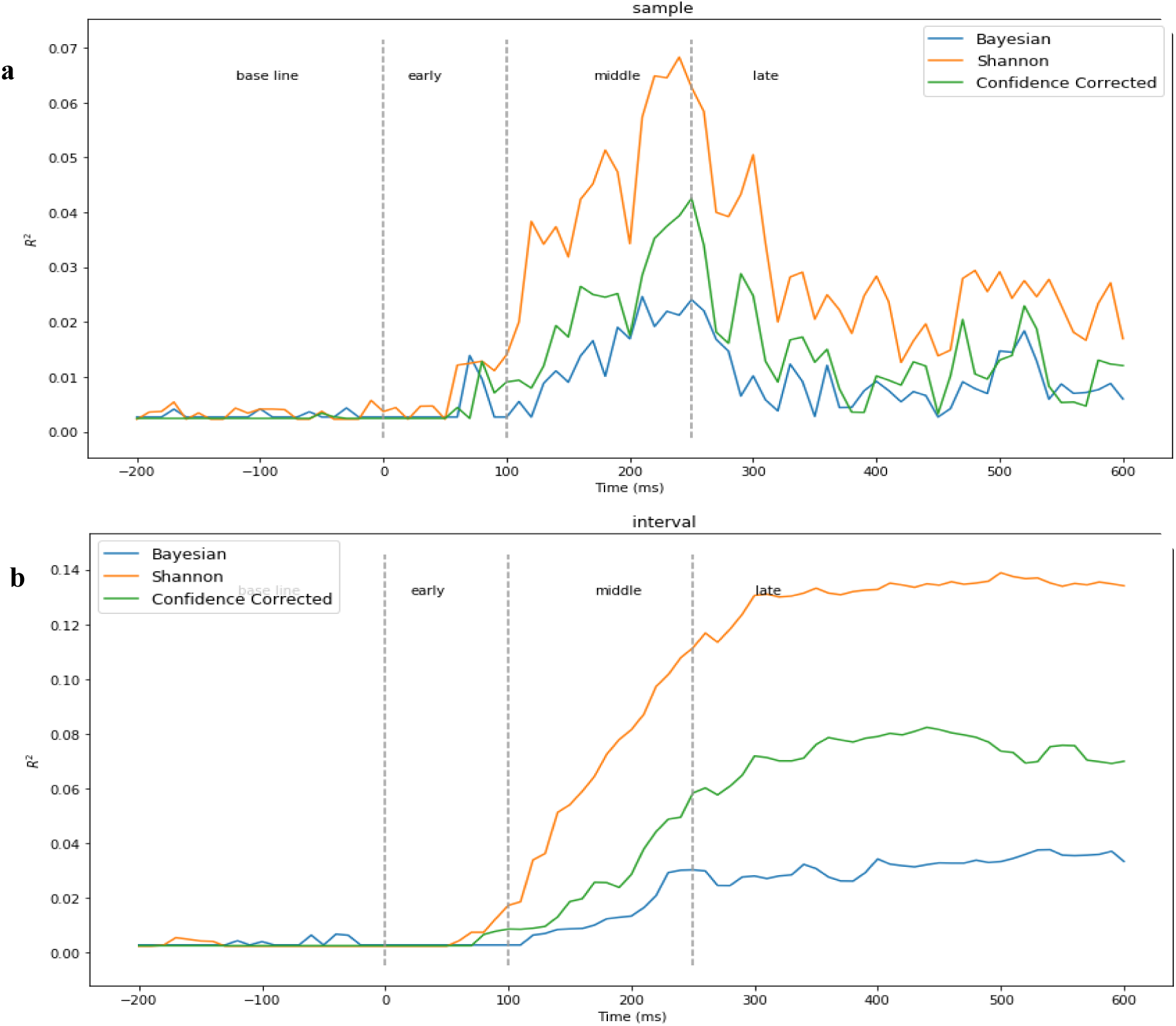
MEG auditory task results. **a)** Surprise decoding powers of the three surprise quantifications for every single temporal *Sample*. **b)** Surprise decoding powers of the three surprise quantifications for the temporal *Interval* [-200ms, t] (colors should be used in printing).

**Fig 4.**
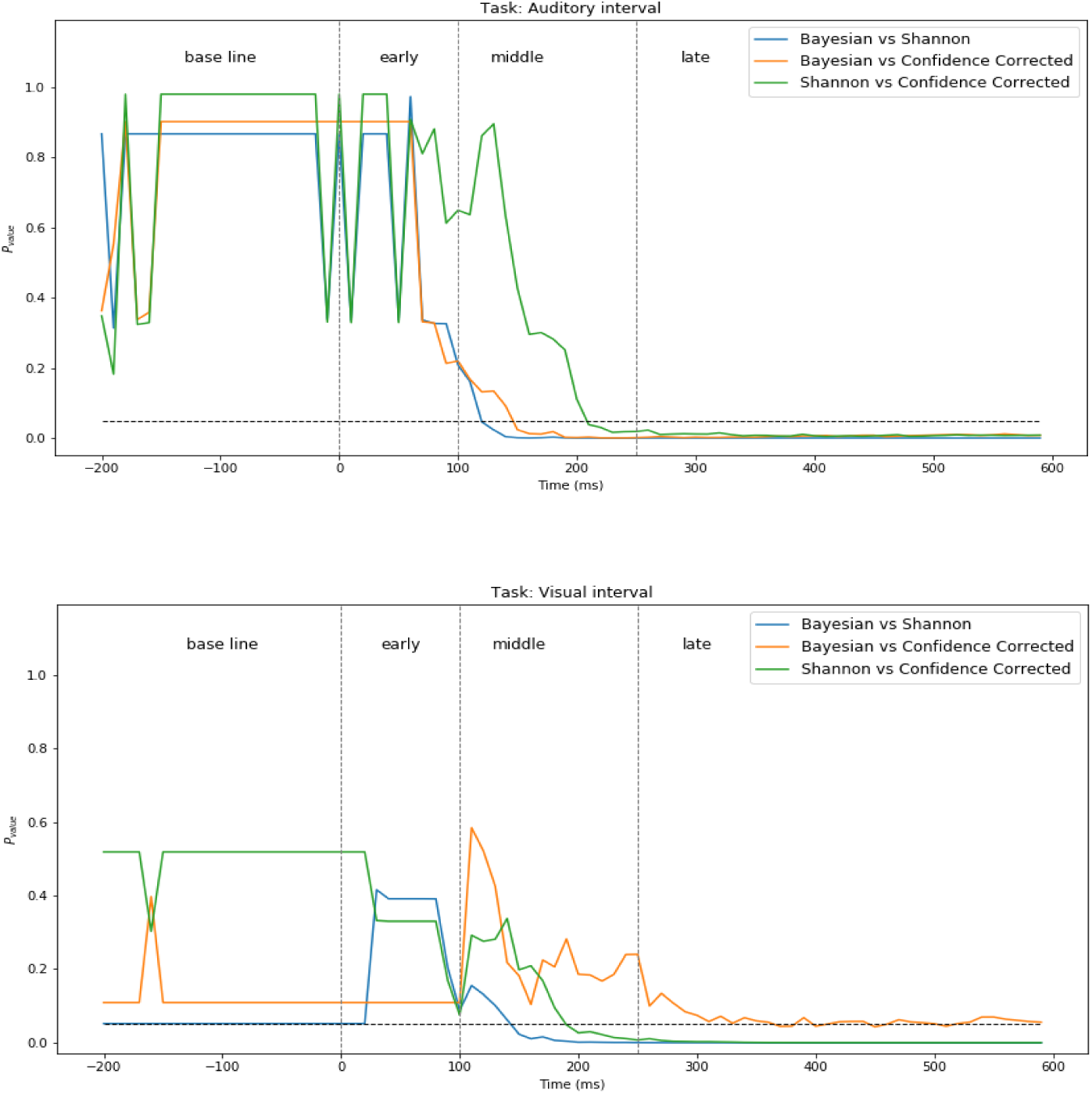
P-values of comparing decoding powers of three surprise quantifications using temporal *Intervals* [-200ms, t] in EEG auditory and visual tasks. Colors should be used in printing.

**Fig 5.**
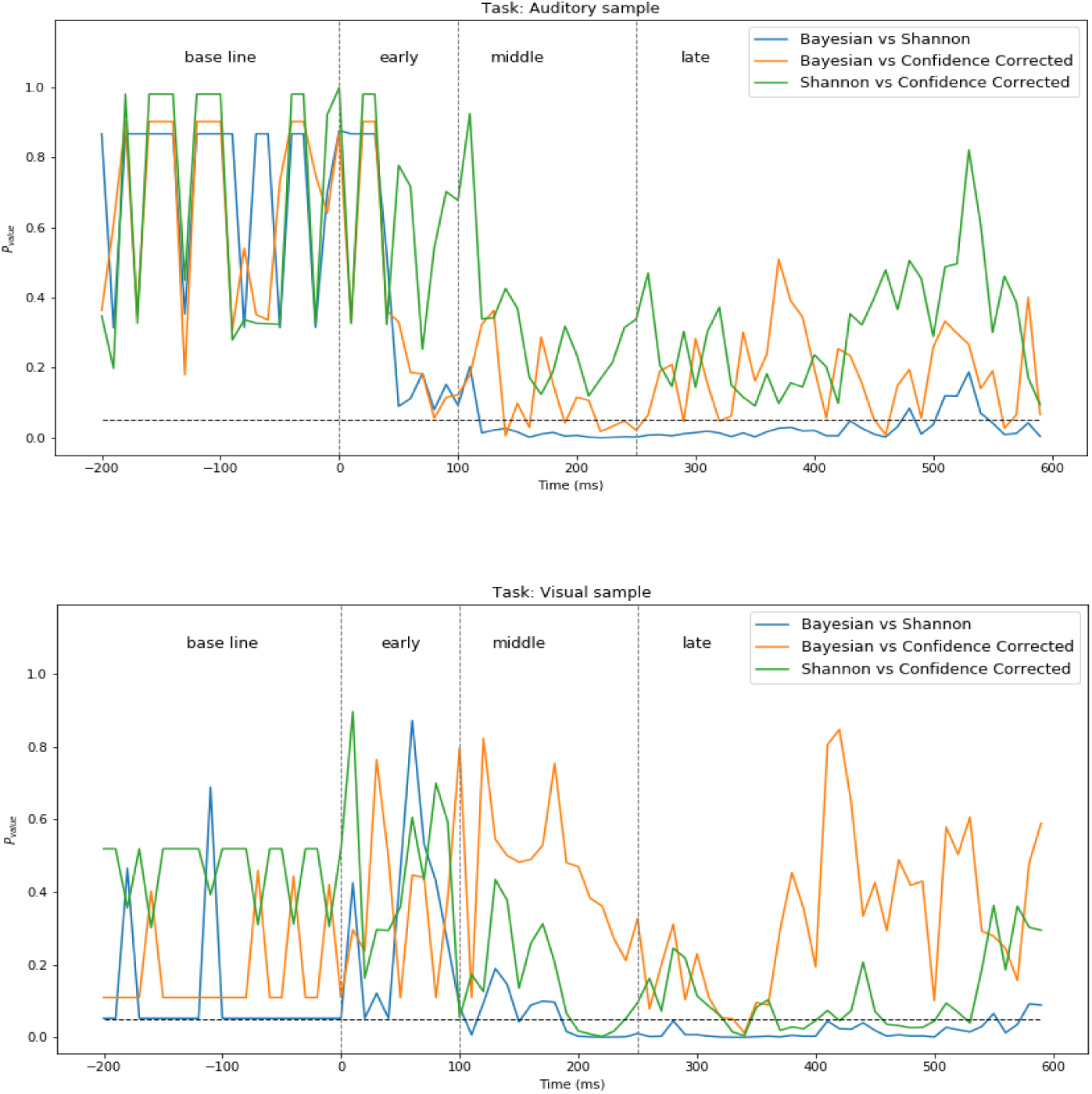
P-values of comparing decoding powers of three surprise quantifications of using each single temporal *Sample* in EEG auditory and visual tasks. Colors should be used in printing.

**Fig 6.**
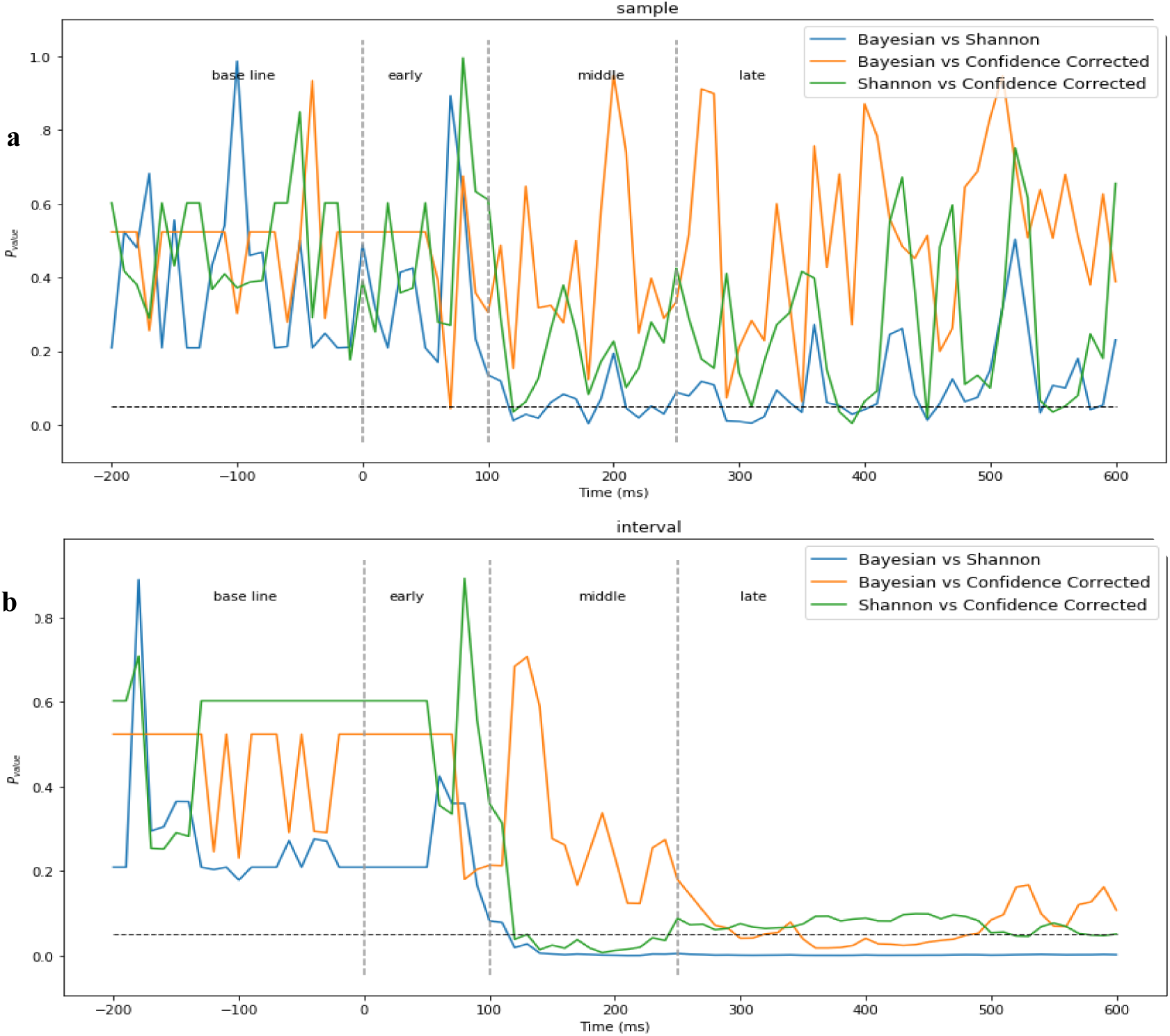
P-values of comparing decoding powers of three surprise quantifications. a) *Sample* and b) *Intervals* regimes in MEG auditory task (colors should be used in printing).

**Fig 7.**
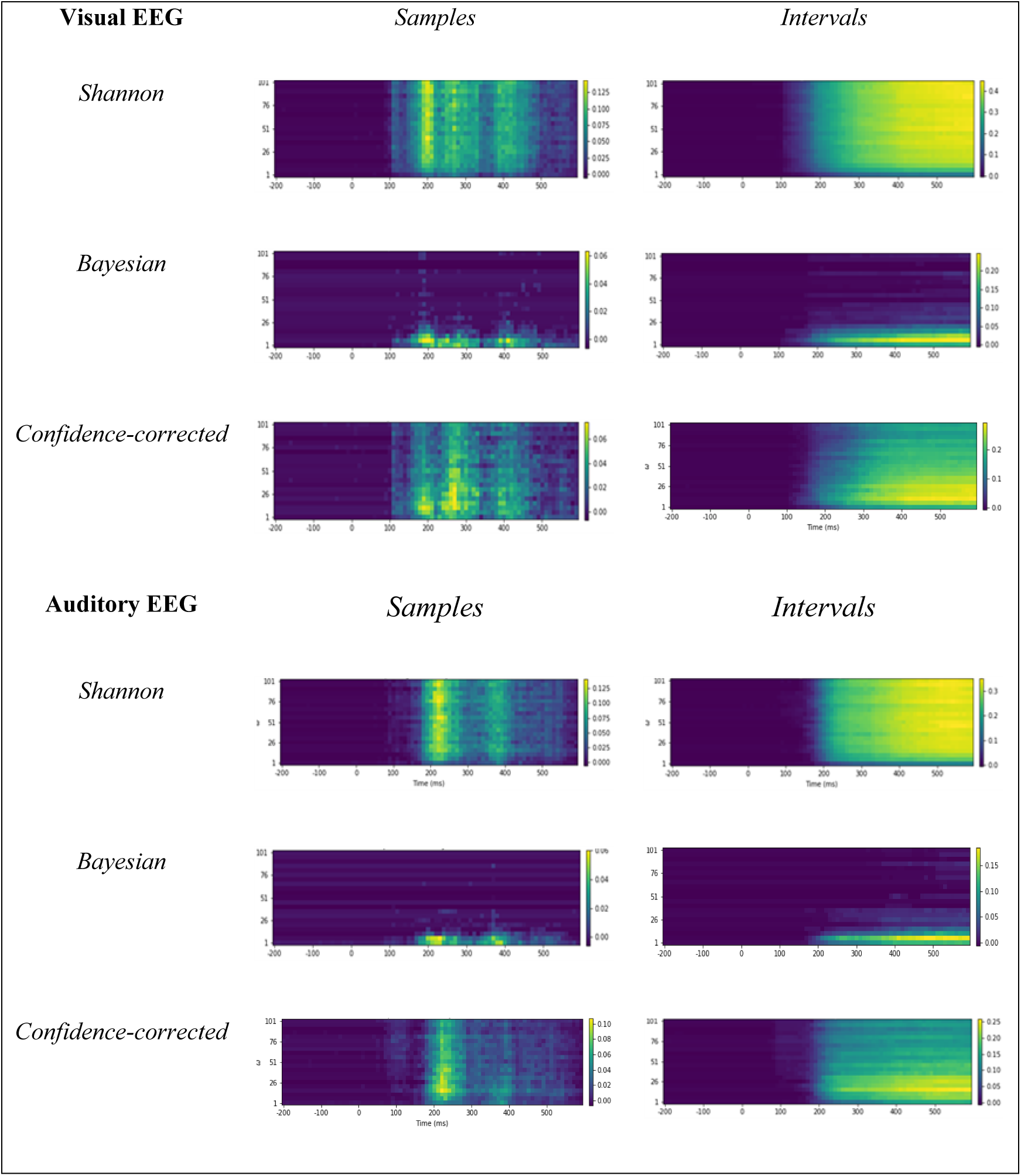
The decoding power of different values of *w* over time for different definitions of surprise, Intervals or Samples regimes and two sensory modalities in EEG data. Colors denote the decoding power and should be used when printing.

**Fig 8.**
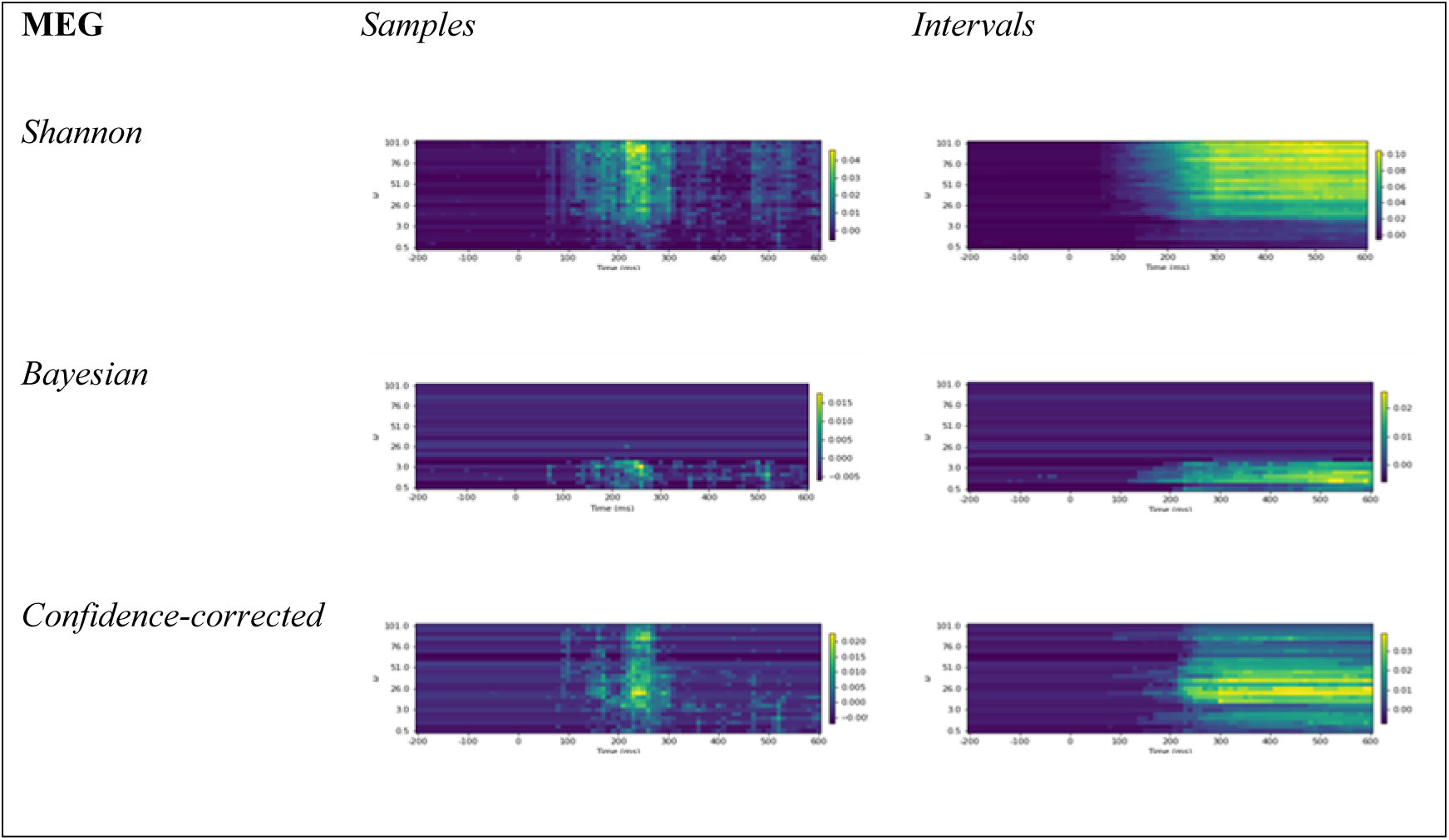
The decoding power of different values of *w* over time for different definitions of surprise, Intervals or Samples regimes and two sensory modalities in MEG data. Colors denote the decoding power and should be used when printing.

### Decoding power

Here we describe the decoding powers of the three quantifications of surprise when employed to supply labels for training surprise decoders. In temporal regimes based on the *Entire epoch* and each of the three *Segments* (of Early, Middle, and Late), there is only one decoder per subject. Hence, the *R*^2^ (R-squared) values that are to be reported will be single numbers for each case averaged on the subjects, which are reported in Table 1. For the temporal regimes of *Interval* and *Samples*, since the target time is swept from the start of the epoch to the current sample time *t*, for each value of *t* there will be a decoder trained, for which an *R*^2^ value will be reported. These *R*^2^ values for these two temporal sampling regimes are illustrated as curves versus time *t* and are plotted in Figs 1 to 4.

#### Entire Epoch and Segments regimes

The *R*^2^ values using the entire samples, as well as the segments named Early, Middle and Late time intervals are presented in Table 1 for both auditory and visual tasks of EEG and the auditory MEG task, and for the three surprise quantifications. The chance level is calculated and shown for every condition to have a criterion for assessing the *R*^2^ values as decoding powers. Comparing the decoding powers achieved in each task and for each definition of surprise with the corresponding chance level, we observe that for the Early segment, the *R*^2^ values and the chance levels are close to each other and hence, the hypothesis of independence between the surprise values (with any surprise definition) and the Early segments is not violated. Unlike this low performance of the Early segment in describing the theoretical surprise values, it can be observed in Table 1 that the Middle and Late segments demonstrate significant values of decoding power.

#### Samples regime

The decoding power of the regression model, whose input feature is a single time sample at a target time *t* of the response in different ICA components, is illustrated in Fig 1 for the EEG and in Fig 3.a for the MEG data, for different values of *t* ∈ [−200ms, 600ms]. Due to the employment of only one time sample in each epoch for decoding the trial’s surprise, it is understandable to have relatively lower *R*^2^ levels. Interestingly, in Fig 1, two major local maxima are observed in each curve. This means that for all definitions of surprise and for both the auditory and visual task data, one can identify one time sample in the Middle segment and one in the Late segment each having a peak decoding power. These time samples with peak *R*^2^ values can serve as statistically meaningful placeholders for the well-known N200 and P300 components of the ERP terminology, respectively. The curves of Fig 3.a appear different from those of Fig 1 in that the second local maximum points in the MEG corresponding to the Late segments are smaller than the peaks of the Middle segment, whereas the maximum points occurring in the Late segments of the EEG analysis in Fig 1 had large values.

#### Interval regime

Figs 2 and 3.b illustrate the decoding powers of decoders trained using an interval of temporal samples in the range of [-200ms, *t*] for different values of *t*. This regime is expected to reveal at which time enough evidence has been accumulated from the response for achieving a confident decoding performance. As expected, all the curves in this regime are approximately monotonically increasing with time as more data is used by the decoder with increasing *t*. In each curve, the *R*^2^ value stays close to zero until around 100ms, when there is a considerable jump in the decoding power. This increase occurs in the temporal range which we called the Middle segment in our *Segments regime*. The decoding power continues to increase gradually afterwards. This behavior is observed for all three definitions of surprise and for both stimuli modalities in the EEG analysis. However, the curves of the MEG and EEG analyses have a difference in that the MEG curves appear more flat after the first rise. This can be explained by the lower decoding power of the Late samples in decoding surprise, which was discussed in the *Samples regime*.

The time point at which the jump in the decoding power occurs in the *Interval regime* is expected to be located on the temporal axis at an earlier time than the point in the *Samples regime* with the highest decoding power. This is because the *Interval regime* tends to accumulate evidence about the surprise from the beginning of the epoch to the present target time, and hence feeds its surprise regressor with a larger amount of information compared to the decoder in *samples* regime, which only uses a single point in time to decide whether a surprise has occurred. Nevertheless, the fact that the *Samples regime* is able to identify one single time point in the Middle segment which can determine whether a surprise has occurred with significance (for any of the three definitions of surprise) is itself a remarkable observation in our study. The gradual increase in the decoding power of the *Interval regime* eventually matches that of the *Entire epoch regime* as more time points are recruited to accumulate evidence for the occurrence of surprise (compare the maximum decoding power of Figs 6 and 7.b with the first row of Table 1).

### Significance of temporal features

Based on the calculated *R*^2^ values achieved by the different temporal samples in decoding surprise, here we discuss the relative significance of these samples or intervals in revealing whether a surprise has occurred.

Table 2 shows the p-values of the t-test for comparing the decoding powers of the three segments of *Early*, *Middle*, and *Late*, employing data from the different subjects as the statistical samples.

For EEG experiments, it can be seen from this Table that there is a statistically significant distinction between the decoding powers of the *Early* and *Middle* segments, and that of the *Early* and *Late* segments (p-values < 0.05). However, we did not observe any significant difference in the *R*^2^ value of the *Middle* and *Late* segments.

For MEG experiments, the Table indicates that there is a statistically significant distinction between the decoding powers of the *Early* and *Middle* segments (p-values < 0.05). However, we did not observe any significant difference in the *R*^2^ value of the *Middle* and *Late* segments and the *R*^2^ value of the *Early* and *Late* segments. This is because Late components in MEG are not as powerful as the ones in EEG data for decoding surprise.

As indicated in Fig 1, for the *Samples regime* which examines the significance of each sample time point in decoding surprise, two main local maxima occur, one in each of the *Middle* and *Late* segments. The first peak always has a higher power in decoding surprise, and its power is noticeably higher in decoding the Shannon surprise. The global maximum of the decoding power curves across time always occurs in the *Middle* segment, reflecting the relative importance of components like N200 in decoding surprise. For the MEG data, Fig 3.a illustrates power of the Middle components in decoding surprise.

In addition, in the monotonically ascending *R*^2^ curves of Figs 2 and 3.b, which illustrate the decoding powers of *Intervals* (i.e. using all temporal samples in a range from the beginning of the epoch to the present target time), a considerable increase in decoding power occurs (only) in the Middle segment, and the curves asymptotically saturate afterwards. This is true for all quantifications of surprise. In fact, these curves indicate that there is a range of temporal samples in the Middle segment which possess a critical role in describing surprise, to the extent that adding samples from later times (e.g. points in the Late segment) does not noticeably enhance the surprise decoding power. In other words, recruiting only the temporal samples in the [100ms, 250ms] interval seems to suffice for describing the theoretical surprise in both the puzzlement and enlightenment senses. To the extent of our knowledge, this observation has not been reported earlier, and furthermore, our analysis is the first to quantify this sufficiency through rigorous statistical analysis and for the different quantifications of surprise.

### Comparison of surprise quantifications

Here we compare the relative powers of the three quantifications of surprise which were employed to create labels for training decoders.

When the *Entire epoch*, or each of the *Segments* of *Middle* or *Late* are used as temporal segments, our results (Table 1) show a clear advantage for the Shannon surprise as the most decodable surprise in terms of the model’s average *R*^2^ value, while the Bayesian surprise is the least decodable surprise. The results for the *Early segment* are not used in this comparison as the decoding powers are close to the chance level, reflecting the very low amount of useful information in this segment for decoding surprise.

Furthermore, for the *Entire epoch*, or the *Middle* or *Late* temporal segments, in the EEG data, the superiority of the Shannon surprise over the other two definitions of surprise is the only significant difference (with p-value less than 0.05) (Table 2). In other words, the statistical difference between the Bayesian and the Confidence-corrected surprises is not significant, while a significant difference exists between the *R*^2^ values of the Shannon surprise and the Bayesian surprise, and those of the Shannon surprise and the Confidence-corrected surprise (Table 3). For a similar temporal segmentation of the epoch for the MEG analysis, the only significant difference occurs between the Shannon surprise and the Bayesian surprise, in favor of the Shannon surprise.

In the case of the *Samples* temporal point selection regime, the Shannon surprise is significantly better decoded than the Bayesian surprise in both visual and auditory tasks of the EEG analysis and in the MEG analysis, while none of the other differences in the *R*^2^ value is significant (Figs 5 and 6.a).

Finally, in the *Intervals* regime of the EEG analysis in which temporal sample points are added progressively, all of the two-way comparisons between the decoding powers of the three surprise definitions are nearly significant (Fig 4), and adding every temporal component does not change the order of *R*^2^ for these three surprise values in the visual task. The only exception is in the visual task in that the distinction between *R*^2^ values of the Bayesian and Confidence-corrected surprise is not statistically significant (Fig 4).

In Fig 6.b, for the *Intervals* regime of the MEG analysis, the only significant difference belongs to comparing the Shannon and Bayesian surprise values.

### The effect of integration coefficient

In this part, we will examine the effect of the integration coefficient *w* on the decoding power of the three quantifications of surprise for the two regimes of

#### Intervals and Samples

In Figs 7 and 8 the decoding powers of the designed decoder are plotted for different integration coefficients in the range of [1,100]. The decoding power is shown by colors defined on the right.

Two local maxima are again observed for the *Samples* regime in the Middle and Late components. In the *Intervals* regime, the decoding power constantly increases in time as expected, and hence the color becomes bolder.

Two different behaviors can be observed for the three surprise quantifications. For the Shannon and Confidence-corrected surprise values, when *w* is not small a relatively high decoding power is observed. However, for the Bayesian surprise *w* needs to be relatively small in order to obtain high decoding powers.

According to the Bayesian definition of surprise, after the brain has observed enough stimuli to correctly learn the generative distribution, there will not be a high amount of surprise in response to any type of stimuli because at that time the difference between the two distributions involved in the calculation of the KL divergence diminishes. In addition, for larger integration coefficients, estimates of the underlying parameter *θ* will be closer to its true value. This improvement enables the Shannon and Confidence-corrected surprises to produce more accurate predictions for the brain surprise

In addition, in the *Samples* regime of the EEG analysis, the best integration coefficient is not much dependent on time. In other words, the best *w* is not much different for the middle and late components (Fig 7).

## Discussion

In this study, we aim to examine how the surprise of an ideal observer is reflected in the temporal data recorded by EEG and MEG systems, and which surprise model can be best described by these temporal measurement samples. On the basis of the Bayesian brain assumption, an ideal observer is described as an observer who attempts to estimate the generative distribution of the incoming stimuli. The estimated distribution is updated after receiving each stimulus. The distribution is parameterized in binary oddball experiments using a transition probability matrix (e.g. as in [13]).

Based on the estimated probability of the stimulus, we have calculated three kinds of surprise values in the present work to predict how surprising receiving each stimulus could be. The three quantifications of surprise employed in our study are: Shannon, Bayesian and Confidence-corrected surprise values.

After calculating the surprise values based on the incoming stimuli sequence, we employed a regression model with a subset of the temporal ERP components as its input features, and one of the surprise values as its labels, to estimate the surprise of the brain’s response. We utilized the decoding model of Modirshanechi, 2019 to extract the surprise of the response from the input feature vector. There is a difference in the decoder we employed and the one introduced by Modirshanechi, 2019. We have not used the second ordinary regression block in [15] to acquire the temporal density of the significant coefficients and instead, the values of the different temporal responses are assessed in our study through the four different defined sampling regimes.

Our results indicate that in both the EEG and MEG data, the Shannon surprise is always significantly better decoded than the Bayesian surprise for all temporal sampling regimes, while it is significantly better decoded than the Confidence-corrected surprise only for the Middle, Late, or Entire temporal segments of the EEG data. As for comparing the temporal parts of the recorded data in decoding surprise, in EEG both the middle and late segments yield high decoding powers not significantly different from each other, while in the MEG data only the Middle components have relatively high decoding powers. Generally, for both data types, the most powerful decoding time instance always resides in the Middle segment, suggesting a good placeholder for the MMN readout. In the EEG data, the best time point in the Late segment, while not as powerful as the best point in the Middle segment, serves as a time point for extracting the P300 amplitude. In the MEG data, the Late components are not powerful enough for decoding surprise. In addition, the results show that in surprise decoding, augmenting the Middle segment with components of the Late segment does not improve the decoding power noticeably. These observations are true for all three definitions of surprise used in this study.

### Evidence for Bayesian brain and ideal observer

In this work, we have constructed our decoding model based on the Bayesian model of the brain by presuming a generative model for the world [39–43]. The brain is assumed to update its perception of the world after receiving inputs according to the Bayesian updating model. Our surprise decoding approach assumes an ideal observer model for the brain, which postulates that the brain attempts to find the distribution from which the input sequence is generated (parameterized by a transition probability matrix), and updates its estimate of this distribution using the Bayes rule [12–15, 27, 29, 38]. Therefore, the assumptions of the Bayesian brain and the ideal observer are embedded in the way we have calculated the theoretical surprise of each stimulus. Although there are three different approaches for defining this surprise, all are based on the parameters learned following the Bayesian brain and ideal observer assumptions.

Our results demonstrate the feasibility of decoding these three quantifications of the theoretical surprise on a trial-by-trial basis with noticeable decoding power, and hence provide new evidence for the Bayesian brain and ideal observer assumptions.

### Finding the best model for surprise

Studying the phenomena associated with the brain’s surprise response is an important aspect of modeling the process of learning in the brain. In these models, surprise has been regarded as a parameter which the brain tries to minimize during learning [14, 16, 17]. One of this paper’s objectives has been to propose an approach to best quantify surprise as a parameter varying with every incoming stimulus that fits the recorded data. Finding the best description of surprise can offer evidence on whether the temporal components of EEG (or MEG) data more strongly reflect the unlikeliness of the input’s value or corroborate the underlying belief updating processes.

The Shannon surprise is a quantification that only considers how unlikely the appearance of a certain input value is. It does not consider the estimated probability of receiving other values for the stimulus. The Bayesian quantification of surprise considers the whole distribution that generates the stimulus and updates estimates of its parameters before and after the arrival of each stimulus.

The Shannon surprise was shown earlier to better correlate with the P300 fluctuations of the response than the Bayesian surprise [12]. Our work demonstrated that the Shannon surprise fits (statistically) significantly better with the data than the Bayesian surprise when the entire epoch or certain segments of the recorded data are used to decode surprise, and that this advantage exists in both auditory and visual sensory modalities in EEG records as well as the auditory modality in MEG data.

The superiority of the Shannon surprise can be explained by noting that the underlying processes associated with the different manifestation of surprise in EEG recording modulate the activity of different brain regions [21–23]. The Shannon surprise has been associated with the activity of the posterior parietal cortex (human lateral intraparietal area), while the Bayesian surprise has been linked to updating processes which occur in the anterior cingulate cortex [22]. As the former activity occurs closer to the brain’s surface beneath the scalp, its presence may be more strongly reflected in the EEG data and hence, be better decoded.

Furthermore, we are using a “Bayesian sampler” approach to describe the behavior of the brain, employing a limited sample size to estimate the parameters of the input distribution [44]. Hence, a possible explanation for the significant difference between the decoding powers of the Shannon and Bayesian surprises might be the effect of limited sample size when estimating the parameters involved in these two models. Further analysis may reveal the particular differences stemming from estimating the mean of a distribution (Shannon model) from a sequence of samples versus comparing an entire distribution to its update (Bayesian update) when a limited number of previous observations is used. This analysis can be the subject of future work related to our results.

### A general consideration of the definition of surprise

It is important to distinguish between the surprise in the recorded response of the brain and the underlying surprise of the sequence of the input stimuli. The former is studied extensively as a parameter to be extracted from the recorded data; for example, in MEG [38, 45], and in EEG, as the amplitude of the P300 component [13, 20, 24, 25, 46], or as the MMN component [27, 28]. The latter terminology for surprise is extracted from the sequence of input stimuli based on an assumption about how the brain estimates the parameters of the input stimuli distribution (for example, the ideal observer assumption used in this work). This surprise is hence not dependent on the actual recorded data from the brain. There are two major approaches for quantifying the surprise of the stimuli which have been reported to be related to the surprise of the brain: the Shannon surprise and the Bayesian surprise.

Aside from these two well-known models for the stimuli surprise, we have also considered a new approach of quantifying the surprise of the stimuli called the Confidence-corrected surprise [14], the correlation of which with the EEG recorded data has not yet been examined (to the best of our knowledge). The dataset we selected for this work contains only two kinds of stimuli; therefore, if an observer considers the probability *p* for the deviant stimuli, the probability of the other stimulus type will be 1 − *p*. Hence, the commitment parameter defined by Faraji, 2018 will not show its importance in our study. Another study with more than two stimuli types might be able to compare the power of this new definition of puzzlement surprise with that of the Shannon surprise.

A remarkable distinction of our work is that we have not considered any single EEG (or MEG) temporal component as a representative for the surprise of the brain. Extracting a reliable value from each epoch (even after epoch averaging, which is a common practice in ERP analysis) is a complex and rather ad-hoc procedure [25, 47–50]. In our approach, we use the entire recorded response to derive the surprise of the brain. Similar approaches to decoding the brain’s surprise have been reported recently in the works of Modirshanechi et al. (2019) and Maheu et al. (2019). In our analysis, for every subject, every sensory modality, and each of the three definitions of theoretical surprise, we have calculated the surprise of the brain as a linear combination of selected components of the recorded data.

The outcome of our data-driven approach provided new evidence for the value of the MMN and P300 components of the EEG epoch as placeholders for the brain’s surprise when decoding any of the three quantifications of surprise. The importance of samples in the Middle and Late components of the ERP response in reflecting the brain’s surprise was established through rigorous statistical analysis of their decoding power for all three definitions of theoretical surprise.

### A general consideration of the location of surprise

Ostwald et al. (2013) postulate that the secondary somatosensory cortex represents the processes associated with the Bayesian surprise. Also, based on fMRI data analysis, O’Reilly et al. (2013) and Schwartenbeck et al. (2016) associate the Shannon and Bayesian surprises with the activities of the posterior parietal cortex and the anterior cingulate cortex, respectively. In addition, the well-known surprise-related components of the ERP signal such as the MMN and P300 have been shown to emanate from the right frontal cortex and the fronto-central regions of the brain, respectively [24, 51]. In the current study, we have not performed any spatial selection among the EEG (or MEG) electrodes or among the ICs with spatial distributions close to the known sources of the different surprise processes in the brain. While this choice offers generality to our analysis through employing all available data and letting the decoders capture all the relevant information during the training procedure, a more targeted approach can also be pursued based on selecting subsets of either the electrode channels or the ICs which are spatially associated with the brain regions known to modulate the Shannon or Bayesian surprise mechanisms. Such an extension to our work might reveal whether the Bayesian theoretical surprise can achieve a higher decoding power than the Shannon surprise in decoding the recorded EEG (or MEG) data in spatially-selected configurations of channels or ICs.

### MMN versus P300 for describing surprise

Earlier studies have used one of the definitions of surprise to describe the trial-by-trial fluctuations of one of the MMN or P300 components [12, 27, 29]. We have considered the role of both of these components with each of the different quantifications of theoretical surprise. We observed that both the Middle and Late components, containing the MMN and P300, respectively, have noticeable surprise-decoding power even when considered separately. However, while we did not observe a significant difference between the decoding powers of the Middle and Late components, our analysis revealed that the Middle components of the ERP can decode the surprise better than the Late components for all three definitions of surprise. As a result, when using only one component of the ERP signal, the MMN value fits better than P300 to describe the stimuli’s theoretical surprise. An extension to this study could be to evaluate whether a decoder trained using the MMN component can efficiently decode the occurrence of a P300 component [52], or vice versa.

### Optimizing the timescale of integration

The timescale of integration, or the integration coefficient denoted by *w* in our model, is a parameter representing the number of recent stimuli values kept effectively (with considerable coefficients) in the memory for estimating the distribution that generates the stimuli [13, 15, 24, 25, 53]. The timescale of integration works between the trials. For each subject and for each of the definitions of surprise, we optimized for the timescale which provided the best decoding power for each of the temporal sampling regimes.

According to the results of our optimization, a large timescale of integration is not suitable for describing the Bayesian surprise. In other words, the best description for the Bayesian surprise derived from the brain’s response occurs when a rather short window of integration is used. This behavior stems from the very definition of the Bayesian surprise, which measures the difference between the distribution estimated for the parameters of interest before and after the arrival of each stimulus. This divergence amount constantly decreases as we increase the timescale of integration since the two distributions become closer to each other.

Given the rather short window of integration involved in keeping track of the Bayesian surprise, this quantification of surprise tends to be more sensitive to the fluctuations of the recorded data compared to the Shannon and Confidence-corrected surprises, which use longer windows of integration, making them more robust to such fluctuations.

Considering the oddball tests used in our study as examples to make this case, if the sequence of stimuli is generated by a random process with fixed statistics (e.g. the probability of the occurrence of a deviant), a longer window of integration allows for estimating the probability of the deviant stimuli with higher accuracy, and hence is optimal for offering a high decoding power.

However, in the case of the Bayesian surprise, on the one hand a long integration window results in diminished divergence in the consecutive updates and hence cannot produce surprise values at par with the brain’s data. On the other hand, a short integration window tends to follow local fluctuations in the statistics and not provide accurate estimates of the underlying statistics of the input sequence generation process. This lack of performance for both long and short timescales of integration may offer an explanation for the inferior decoding power of the Bayesian surprise across all of our experiments.

### Motor response effect

In this work, we have studied the EEG recordings with a motor response by the subjects to the deviant stimuli. In order to examine whether the motor response of the participants had a confound effect on the decoding result, here we analyze another auditory EEG dataset which includes no motor responses. This analysis allows us to evaluate the power of our decoder when motor responses are not part of the oddball task. Earlier, the work of Modirshanechi et al., 2019 reported no significant difference between the decoding powers of pairs of oddball EEG recordings as follows: 1) Between the decoding power of the dataset with asymmetric motor response (the EEG dataset used in our study in which subjects pressed a button only in response to the deviant stimuli) and the decoding power of another EEG dataset with symmetric motor responses (in which the subjects pressed a button for both the standard and deviant stimuli); 2) Between the decoding powers of the dataset with asymmetric motor response and another EEG dataset with no motor response. Despite this observation, it is still probable that the strong decoding power of the late components of the EEG data may be due to the motor responses even though the surprise can still be decoded based on the late components that do not have any motor response confound.

To address this, we applied our decoding methodology to the EEG dataset with no motor response in the *Samples* regime to evaluate the decoding power of the Late components in comparison to that of the Middle components.

This dataset includes four different auditory oddball tasks and is openly accessible at www.bnci-horizon-2020.eu/database/data-sets with the name “auditory oddball paradigm during hypnosis”. There are data from two healthy subjects, one female and one male, in this dataset. Two of the tasks are performed in normal awake condition, and during the other two tasks the subjects were hypnotized. We used the data in which the subjects passively listen to the auditory stimuli.

The subjects were presented a random sequence of 420 short complex high tones (440 + 880 + 1760 Hz) and 60 short complex low tones (247 + 494 + 988 Hz). The duration of each stimulus was 50 ms. The EEG data were recorded with a sampling rate of 512 Hz using active electrodes in 27 channels and two pairs of electrodes recorded EOG (Electrooculography).

The steps of EEG preprocessing described in Materials and Methods were similarly applied of this dataset. The dimension of each feature vector was *p* = 80 × 27 = 2160 (80 samples in every epoch multiplied by the number of channels), and after removing noisy epochs approximately *N* = 440 features remained.

Fig 9 depicts the decoding power of each temporal sample attained using our surprise decoding method regressed the Shannon surprise. It can be observed that the Late component responses still possess a strong decoding power comparable to the decoding power of the Middle component responses. It can be hence deducted that the relatively strong decoding power of the Late components of the main EEG dataset in our analysis plotted in Fig 1 is not due to the motor response confound.

**Fig 9.**
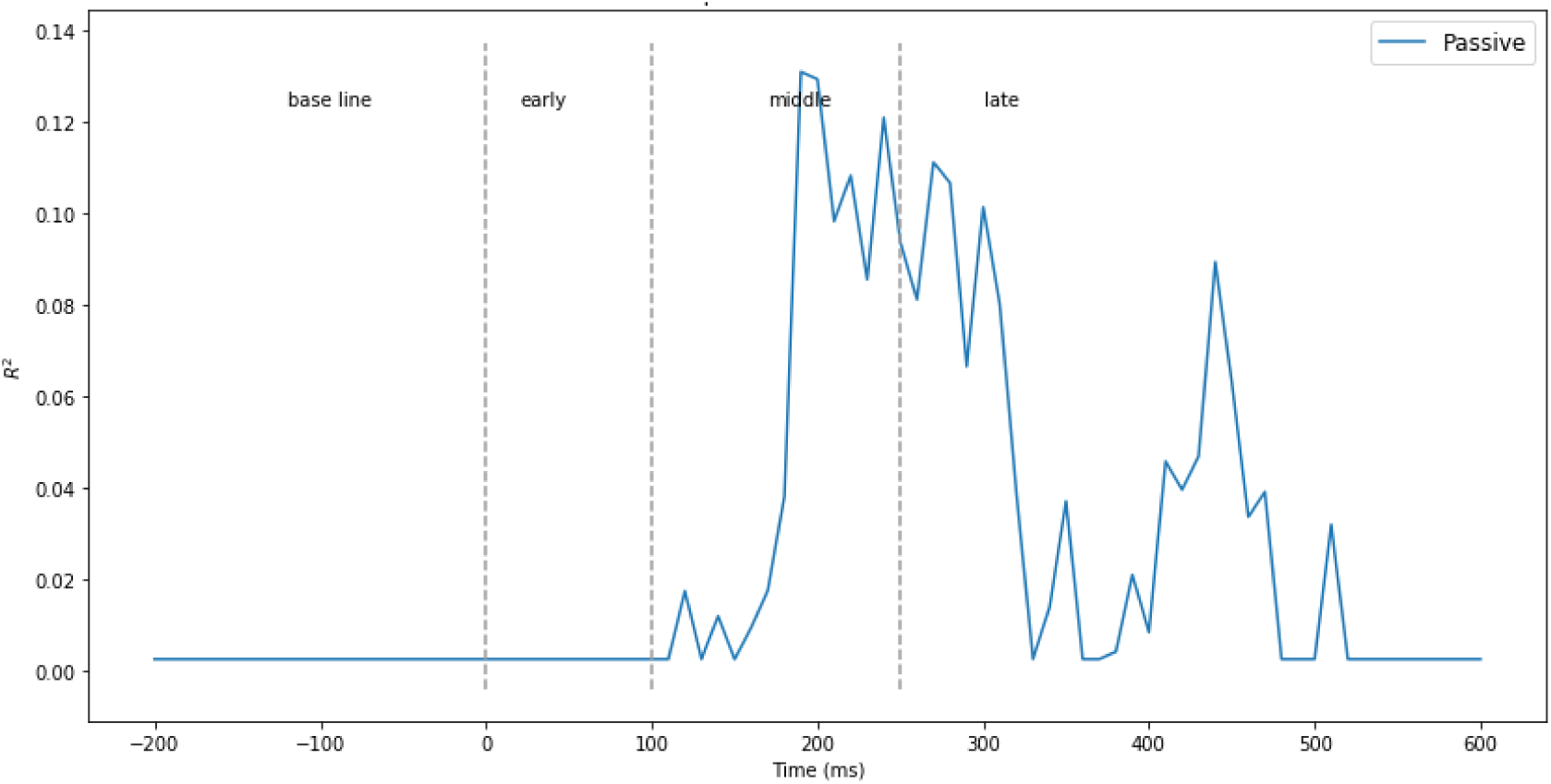
The power of every single temporal *Sample* for decoding Shannon Surprise in both active and passive tasks of benchmarking EEG dataset.

### MEG and EEG comparison

Better signal-to-noise ratio and readability of the MEG recording compared to EEG [35, 54] offer opportunities for further examination of the mechanisms that generate surprise in the brain. A larger number of recording sensors distributed more densely across the head provides better coverage of local activity beneath the scalp. In our study, we observed that the Late components of the MEG epoch had noticeably less decoding powers than the Middle components. This was a distinctly different behavior from the EEG analysis results in which both the Middle and Late components of an epoch offered significant decoding powers. A likely explanation for the lower performance of the Late components in the MEG analysis can be that while each EEG sensor collects and integrates data from a rather distributed set of sources in the brain, each MEG sensor can only capture the activities of sources in its close proximity beneath the scalp. The surprise generation mechanisms of the brain transmit signals to a number of different regions of the brain, which in turn produce the Late components of surprise which are distributed and diffused. Since these Late components are generated by distributed sources, MEG sensors may not be able to adequately capture them [34, 55].

### Limitations and extensions

Maheu et al. (2019) argue that the early brain responses to an oddball experiment are best explained by the learning of the deviant-to-standard frequency, while the late responses are best explained by the learning of transition probabilities between the two input types. They further propose the presence of both long-term and short-term sequence learning mechanisms in the brain, with the long-term learning process resulting in the early to middle responses observed in an oddball experiment while the short-term learning process being active during the late stage response period [38].

In the datasets analyzed in our work, the probability of the occurrence of the deviants is fixed and small (1/5 in the EEG datasets and 1/3 in the MEG dataset). This probability, called the item frequency in some literature [13, 38], along with a second statistical parameter, the alternation probability, fully determine the characteristics of the input sequence generation. To assess the performance of the two-parameter decoding models discussed earlier, experiments in which these two probabilities take different values would be needed. In addition, the decoding model based on the set of two transition probabilities between the two stimuli types can be analyzed using such experiments. The datasets analyzed in our work did not include variations of the statistical parameters and hence allowed for only a partial study on the role of the sequence generation parameters. As a result, we have not compared the decoding power of calculating surprise based on the item frequency with the one based on transition probabilities as in [13, 38]. In order to analyze models based on both the simple parameters like the item frequency as well as models utilizing higher-level parameters like transition probability matrix, a more diversified dataset with higher probabilities for the deviants, and including various alternation and repetitions probabilities would be required. One might analyze the other three blocks of MEG data that was used in this paper to be able to compare the two ideal observers estimating transition probability matrix and item frequency [38]. Furthermore, analyzing surprise on a task including more than two types of stimuli (like the tasks reported in Mars et al. (2008) and Seer et al. (2016)), with parameters more complex than a mere item frequency, can offer new insight into understanding the mechanisms of surprise in the brain.

The decoder utilized in this paper was proposed by Modirshanechi et al. (2019) based on using single-trial EEG (or MEG) data to regress the surprise of the stimuli. To train the decoder, each epoch of the EEG (or MEG) data is used without considering its temporal order in the experiment. As such, the effect of experience by the participant during the presentation of each block of trials was ignored in our analysis. In an extended analysis, one may consider the order of the current stimulus in selecting the best definition of surprise and extracting the temporal components with highest decoding powers. Such analysis may reveal advantages for the Bayesian quantification of surprise by taking into account the learning process that occurs during the experiment, as the brain learns the generative distribution of the incoming sequence of stimuli.

## Materials and methods

Fig 10 provides an overview of the overall flow of data and the decoding approach used in our analysis.

**Fig 10.**
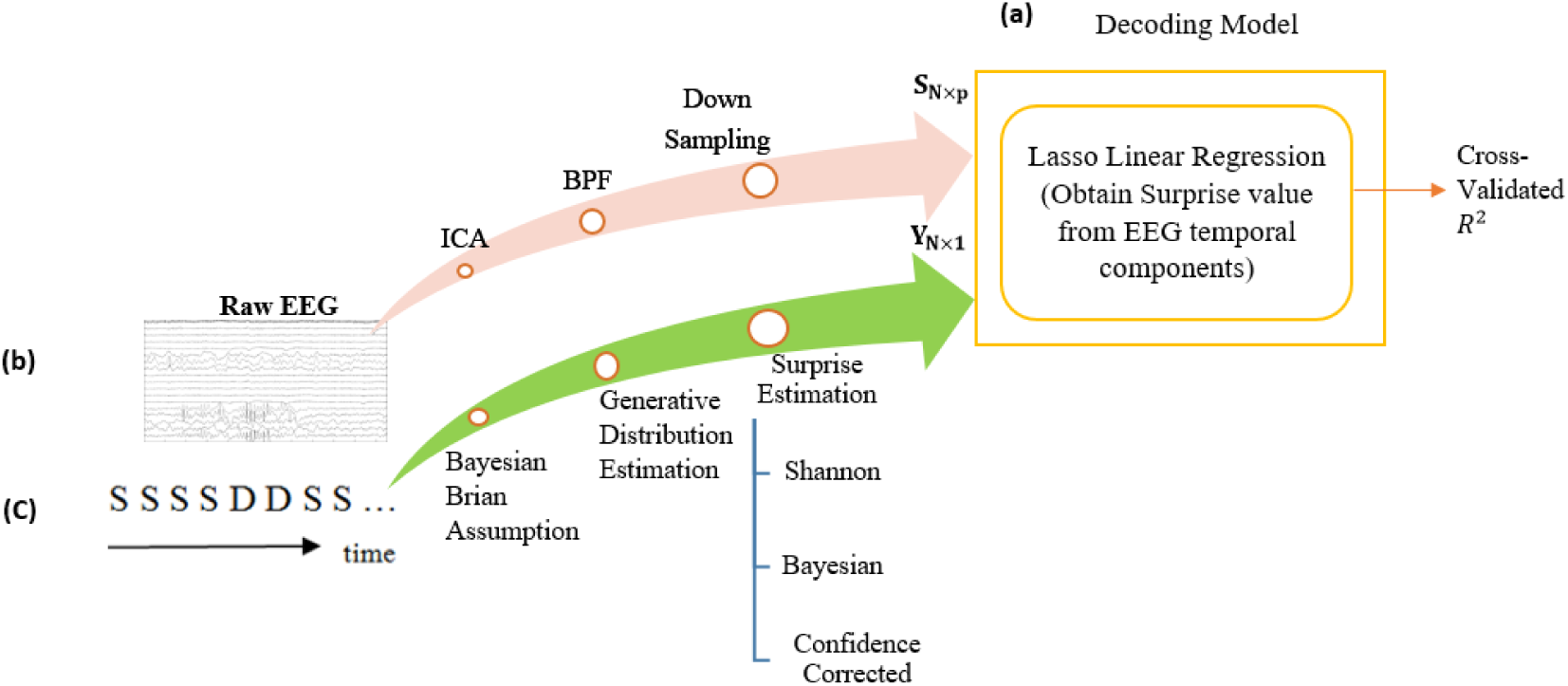
The overall diagram of the decoding model. **a)** The scheme of the decoding model and machine learning tools. **b)** The processes performed on the recorded raw EEG to acquire features for regression. **c)** Surprise calculation module using the oddball sequence of stimuli as input and generating labels for training the regression model.

### Dataset and task

#### EEG dataset

Our analysis is applied to two EEG oddball test datasets with visual and auditory tasks. Each dataset contains simultaneously recorded EEG and fMRI signals and has been published [56–58]. We applied our analysis to the EEG signals of both the auditory and visual datasets. For further clarity and in order to better motivate our results, the unique features of these datasets relevant to our study are briefly described below.

In both sensory modalities, the rare stimuli occurred with a probability of 0.2 and the repeated stimuli occurred with a probability of 0.8 in three different runs for each participant, each run consisting of a sequence of approximately 125 stimuli. Each stimulus lasted about 200 milliseconds. The interval between two consecutive stimuli had a uniform distribution between 2 and 3 seconds. The visual stimuli types consisted of a small green circle as the standard and a large red circle as the deviant. In the auditory mode, the standards were single tones of 390 Hz and the deviants were generated as a broadband “laser gun” sound. The first two stimuli of each run were always fixed to be standards.

The EEG signal was sampled at a frequency of 1000 Hz by a conventional MRI-compatible EEG system using bipolar electrode pairs in 49 different channels, re-referenced to a 34-electrode space. The precision of recording was 1*μ*V.

○ There were 17 participating adults in the experiments with labeled age and gender (six females, eleven males; age mean: 27.7 years, age range: 20-40 years). All participants took part in both auditory and visual tasks. They were asked to press a certain button only at the time of observing deviants and their response latency was also recorded.

#### MEG dataset

Our study is also applied to the recorded MEG data of an auditory oddball task recently published by [38] et al., 2019. In this dataset MEG was recorded for oddball tasks conducted with four different probabilistic models of stimuli generation. We selected the frequency-biased model for stimuli sequence generation, for which the statistical model of the resulting oddball sequence is similar to that of the stimuli sequence of the EEG dataset. In the MEG task, the standard and deviant stimuli are two different French syllables randomly drawn from a binomial distribution with the probability of the frequent syllable being 2/3 and that of the deviant syllable being 1/3. Each syllable lasts about 200 milliseconds and the interval between two successive stimuli is 1400 milliseconds. Each block consists of 400 to 409 stimuli.

Participants include 11 females and 9 males, aged between 18 and 25. The data of two subjects were removed because of their excessive head movements. To ensure that the participants paid attention to the task, they were asked every 12 to 18sounds to predict the next stimuli (being a standard or a deviant) using one of two buttons.

The brain activity was recorded by 306 channels (102 magnetometers and 204 gradiometers) with whole-head Elekta Neuromag MEG system using a sampling rate of 1 kHz and a hardware-based bandpass filter of 0.1 to 330 Hz. Raw MEG data was corrected for head movement and bad channels, and was also cleaned from powerline and muscle movement artifacts. Then, a low pass filter below 30 Hz and a 250 Hz down-sampler was applied to the data. Eye blinks and cardiac artifacts were removed using ICA (Independent Component Analysis). Finally, the data was baseline corrected using a window of 250ms before the stimulus onset [38].

### Preprocessing

#### EEG preparation

The raw EEG signal wasprocessed in our analysis through the following steps (Fig 10.b):

In the first step, the InfoMax ICA algorithm [59] was applied to the data of each subject using the EEGLAB toolbox [60]. Our aim was neither the rejection of artifacts nor selecting among the resulting ICA components. Instead, we assumed a general model and utilized ICA just in order to separate different independent sources of the EEG signal, and make the features of the subsequent surprise regression model as independent from each other as possible. We anticipated that the regression coefficients of surprise-irrelevant components would be zero, and hence a sparse model would be a suitable fit as described further below. Data of each participant for each sensory modality was transformed from the electrode space to the independent component (IC) space by the ICA method. The EEG signals were re-referenced to 34 electrodes, so the resulting IC space was also of dimension 34.

In the second step, a bandpass filter [0.5-38 Hz] was applied to the transformed EEG signals to eliminate undesired frequency components. Then, considering an 800ms period for the acquired EEG data in each epoch, consisting of 200ms baseline before the stimulus onset and 600ms after the onset, subsampling by a factor of 10 produced a sample vector of 80 points for each electrode in each epoch (i.e. a trial in the oddball terminology). Then, the sample vectors of all the 34 electrodes were concatenated to construct a feature vector with the length of *p* = 80 × 34 = 2720 for each epoch. After removing noisy epochs, around *N* = 360 epochs (from a total of nearly 3 × 125 = 375 epochs) were left to be used for training a decoder in the subsequent steps of the analysis. Therefore, the input of our regression model would be a *N* × *p* feature matrix denoted by *S_N × p_* (Fig 11).

**Fig 11.**
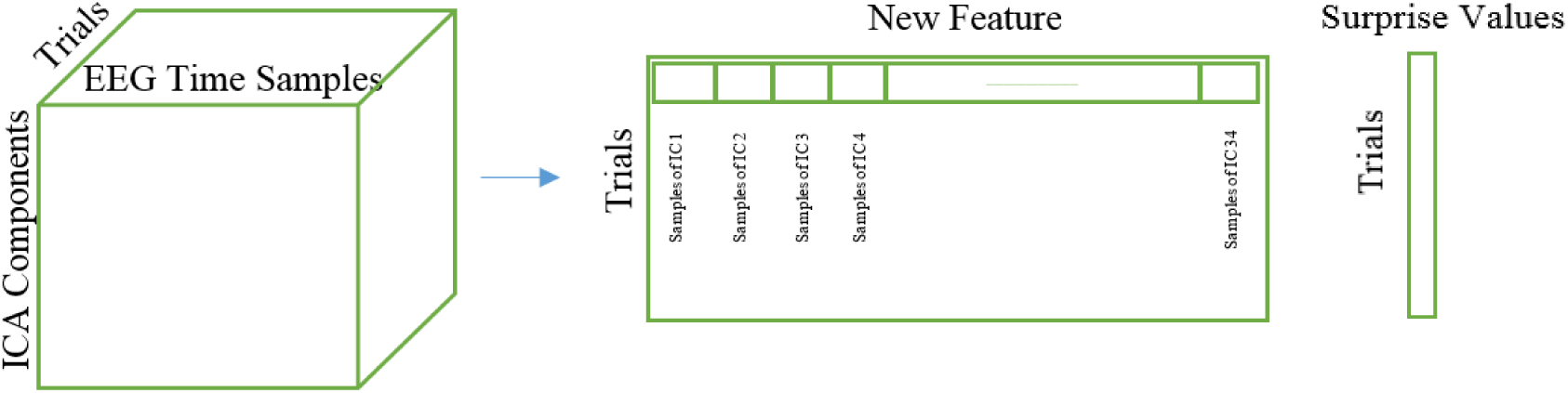
The feature matrix and the surprise label vector used to train the decoding machine.

As will be described later, we defined four different regimes of temporal component selection within each epoch in order to assess the relative values of various time points or temporal segments in the EEG epoch in offering surprise decoding power in the decoding model we used. Hence, not all the 80 time points defined above as feature vector were used in the different regimes, and *p*, the dimension of the feature vector, varied between the four regimes.

#### MEG preparation

The MEG data was initially preprocessed by its developer for head movement correction, bandpass filtering, down sampling and ICA-based artifact rejection [38]. However, in order to employ independent sources as features of the regression model, we performed ICA analysis using the entire 306 channels of the MEG data using the FastICA algorithm [61]. We have chosen FastICA for this data because of the high number of channels (306) which could render the InfoMax algorithm excessively slow. We ended up with an average of 69 independent components for the whole sensors using FastICA. Although in some earlier studies [34, 38] the analysis is done separately on the data of the magnetometers and gradiometers of the MEG recording, we have run the ICA method on the entire 306 sensor set composed of 102 magnetometers and 204 gradiometers in order to use all the available information and to take advantage of the two types of sensors that record different directions of the magnetic field and the spatial gradients [36]. We also considered the interval of [-200ms, 600ms] as the response period and reduced the number of samples by downsampling to 80. Therefore, for the MEG data, the number of features used for training was *N* ∈ [400, 409] (equal to the number of stimuli, which varied between the participants), and the dimension of every feature was around *p* = 80 × 69 = 5520 (equal to the number of time samples multiplied by the number of independent components).

### Ideal observer model

A fundamental question in the Bayesian brain literature is how the brain learns the distribution of the sensory stimuli. The brain is assumed a near-optimal estimator for the probability of the input sequence based on a generative model with Bayesian inference [12, 13, 15, 19, 62, 63]. To be more precise, the brain uses prior belief about the environment, and updates it after each stimulus arrives. In addition, in order to initialize the inference process, it is presumed that the brain begins with the assumption of equally probable input types despite exposure to any possible previous blocks of stimuli [11, 13, 64, 65].

Here, two crucial questions are what exactly constitutes the statistics that the brain attempts to learn from the recent history of observations, and what mechanism is employed by it to arrive at an optimal estimate of this probability. Rubin at al. (2016) proposed that in order to have maximum predictive power for the future stimuli while maintaining a certain complexity level, the best single-parameter model would be to describe the past sequence of stimuli with the number of occurrences for every stimulus type. For example, in a binary oddball paradigm task, the number of standard (or deviant) stimuli up to a certain time is a sufficient statistics deduced as a single-parameter representation of the past for making an estimate for the probability of the next stimulus being a standard or a deviant [19].

#### Transition probabilities

In an oddball experiment, every stimulus can be denoted by a binary random variable *x^i^* for *i* = 1,…,*T*, where *T* is the length of the stimuli sequence. We consider *x^i^* = 0 if the *i^th^* stimulus is standard and *x^i^* = 1 otherwise. This variable follows a Binomial distribution with parameters *p*_0_ and *p*_1_ = 1 − *p_o_* as the probabilities of the standard and deviant stimuli, respectively. Based on the hypothesis that the sequence of items has been generated by a “Markovian” generative process, the sequence can be modeled by the probabilities of transition between the stimuli types. For a binary oddball sequence, the transition probabilities can be stated as a 2 × 2 matrix, which can be estimated by counting the number of successive transitions [13]. It has been demonstrated that utilizing the transition probability matrix for describing the stimuli sequence will statistically outperform the single-parameter approach to describe the brain’s response [13]. For a binary oddball sequence, the definition of the model parameter *θ* can be stated in the form of a 2 × 2 matrix:

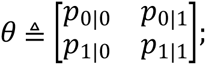

where *p_a|b_* is the probability of transition from stimulus type *a* to stimulus type *b*. Since the sum of each column of this matrix is equal to 1, we can reduce the model parameter’s definition to a vector ***θ***:

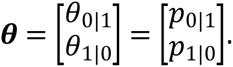

Based on this definition, the likelihood of a sequence of observations ***X****^j^* with length *j* will be:

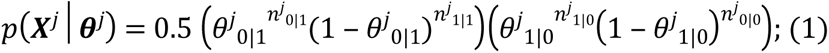

where

***θ****^j^* with elements *θ^j^*_0|1_ and *θ^j^*_1|0_ is the estimated parameter vector after receiving *j* inputs denoted by the vector ***X****^j^*, the probability of the first stimulus is assumed to be ½, and *n^j^_a|b_* is the number of transitions from stimulus type *a* to stimulus type *b* in the *j* observations up to the present sample.

The parameter *n^j^_a|b_* can be computed in different ways [13]. In this paper we have selected the leaky integration method to account for earlier observations. In this method, the most recent stimulus is given a maximum weight and the weights of the preceding observations decrease exponentially with a parameter *w* (the integration coefficient) moving backwards toward earlier observations.

Eq. (1) is the product of two Binomial distributions, each representing one of the two elements of the vector ***θ***. Using the Beta distribution notation to represent the prior probability of these elements as the conjugate prior of Binary distribution, the posterior distribution of ***θ****^j^* after *j* inputs will be the multiple of two new Beta distributions:

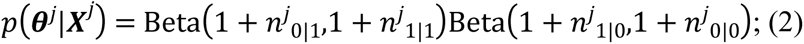

To sum up, the posterior probability of the stimulus-generating Binomial distribution parameter is obtained with a two-dimensional descriptor parameter in Eq. (2). The next step is to use this equation to calculate the theoretical surprise inherent in the stimuli sequence.

### Surprise definition and calculation

Our analysis compares three definitions of surprise for their suitability in modeling the brain’s response in binary oddball experiments. In our decoding model, surprise labels *Y_N_* _× 1_ are employed as labels in the process of training the decoder (Fig 10.c, and 12).

**Fig 12.**
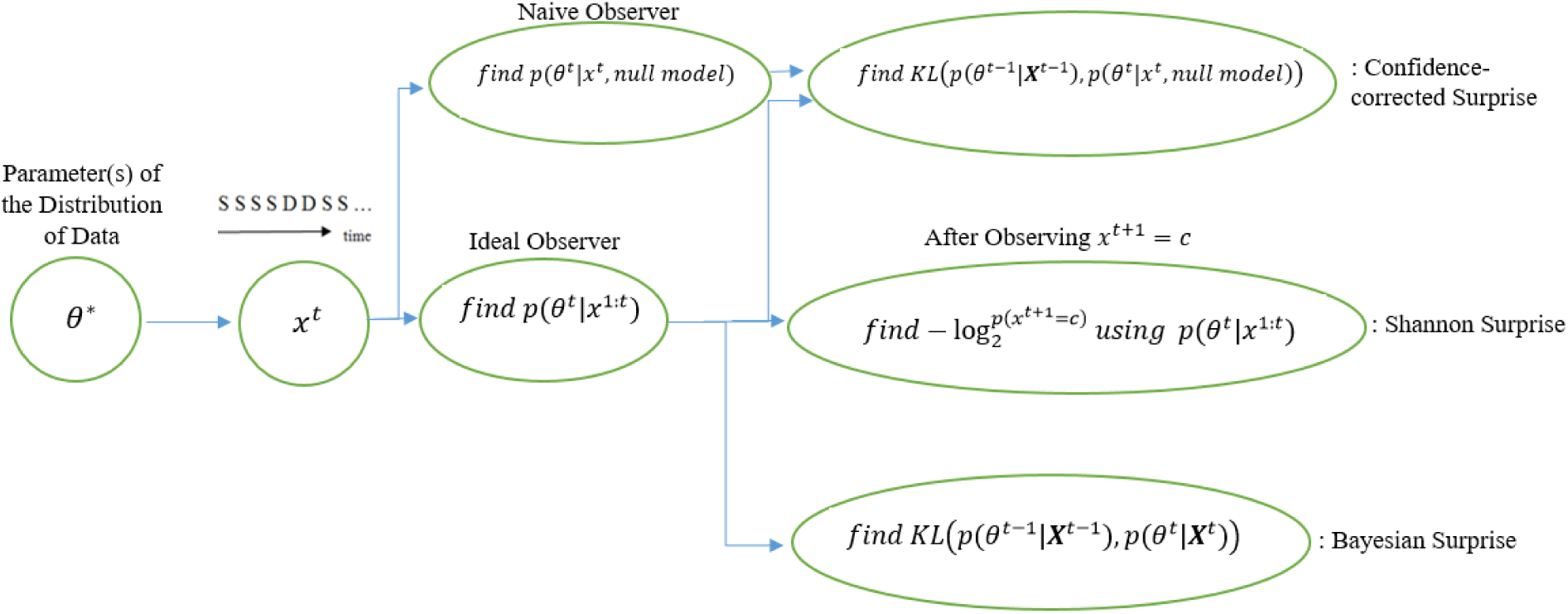
Calculating the three different quantifications of surprise.

Estimation of the stimulus-generating distribution leads to a prediction about the next stimuli, which if violated may produce a “surprise” response by the brain [12, 13, 15, 27] reflecting the prediction error [19, 27].

#### Shannon surprise

The Shannon surprise provides a measure of surprise based on the predicted probability for an observed event [10, 11, 13, 15]. The more unlikely an observation is, the more surprising it will be. The Shannon surprise can be computed for the case of the *j^th^* stimulus having a value *c*:

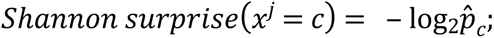

where c is 0 or 1 in an oddball test and 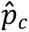 is the predicted probability of stimulus *j* to be *c* and is obtained using this formula:

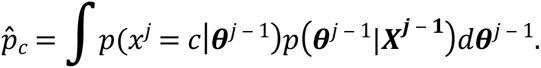

For the single-parameter definition of *θ* (i.e. when *θ* is chosen as the item frequency), we have:

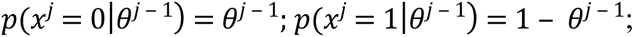

where *θ^j − 1^* is the estimated value of *θ* at the preceding sample. In the two-parameter model, ***θ*** is a vector of the two probabilities of transition. It can be shown that the probability of the *j^th^* item having a value of 0 or 1 can be estimated using the estimated value of ***θ*** at the preceding time (i.e. 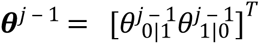) via:

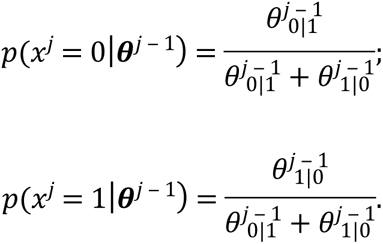

#### Bayesian surprise

It measures how much difference is made in the estimated generative distribution after observing every stimulus. In fact, the surprise can be calculated using Kullback–Leibler (KL) divergence of two distributions assessed before and after observing every observation [66, 67]. For two-dimensional *θ* parameter, the surprise after observing *j^th^* stimulus is:

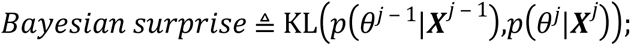

which is the distance between the distributions estimated exactly before stimulus j, having j-1 observations, and after it.

#### Confidence-corrected surprise

Assume a choice reaction time task, with four types of stimuli [*A*,*B*,*C*, *D*], created two models of the world for two observers. The first observer believes that all stimuli types are equally likely (i.e. the probabilities of the four types are [0.25, 0.25, 0.25, 0.25]), while the second observer estimated the distribution as [0.75, 0.25, 0, 0]. After receiving an input *B*, both observers will have the same puzzlement surprise equal to 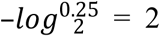 in the Shannon’s model of surprise. But it is obvious that the first person is surprised less than the second one due to the indifference in the predicted probability of occurrence for all types. In other words, the first person has no commitment to the world.

The most recent definition for surprise, calculating it in the puzzlement sense, considers how observers commit to their belief about the world, and attempts to address the above issue [14]. It measures the distance between the distributions estimated for model parameters after observing all the *j* − 1 samples, and the posterior belief of a naїve observer for parameters after observing the *j^th^* sample, *x^j^*. The latter is the posterior probability of parameters considering a uniform prior, after receiving *x^j^* as the first observation. Therefore,

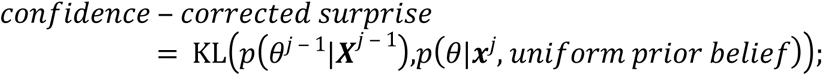

where *θ* can be a number or a vector. For the binary oddball paradigm case, this can be modeled as:

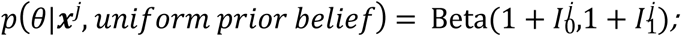

where 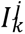 is the identity function (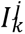 is one if *x^j^* = *k*, and zero otherwise) in the single-parameter model. In the two-parameter transition matrix model:

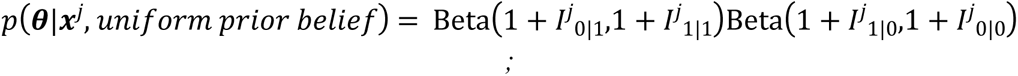

with 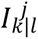 defined as one if *x^j^* = *k* and *x^j^*^−1^ = *l*, and zero otherwise.

Each of these three definitions of surprise yields a vector *Y_N_* _×1_, with elements to be used as surprise labels for each of the *N* trials in training a surprise decoder. The results of using these definitions of surprise will be compared in terms of their ability to characterize trial-by-trial fluctuations of the EEG record after a decoder is trained (Fig 10.c). The analysis is performed for a binary oddball paradigm task in terms of decoding surprise responses, and comparisons between the predictive powers of the surprise definitions are made based on the two-parameter model described earlier.

### EEG temporal samples and segments

In each epoch of the oddball task, four basic time intervals are often identified for coarse-level segmentation of the response profile: From −200ms pre-stimulus to time 0 (Baseline), from time 0 to 100ms (Early Components), from 100ms to 250ms (Middle Components), and from 250ms to 600ms (Late Components). In our study, we seek to identify the significant single time instances or time intervals which can best regress the surprise level. Hence, we define four different regimes of sample selection from the temporal data record, and apply our decoder operation in each regime to find out the most significant set of points in time which can be used to predict the surprise level of the input stimuli (Fig 13). The four regimes of temporal data selection are described below. Note that the entire set of data covering one epoch is 80 points (see “Preprocessing of Data”), and there are 34 channels of data after the ICA function is applied to the electrode signals.

- **Entire epoch:** The total response time (−200ms to 600ms) is used for regression to identify all significant coefficients..
- **Segments (Baseline, Early, Middle, and Late):** To evaluate the regression power of the target time interval, only a pre-defined range (for example late components) is used as input feature vector to the decoder.
- **Intervals:** To evaluate the significance of an interval of accumulated temporal samples from the beginning of the epoch to the current target time, the interval of −200 to time *t* is used as input to the decoder. This operation is repeated for all values of *t*.
- **Samples**: A single sample at time *t* is regarded as the decoder’s input, and this operation is repeated for all values of *t*. Our study applies each of these four temporal sample selection regimes to three decoders, each using one of the three definitions of surprise described earlier. This is done independently for each subject and for each of the two input modalities.

**Fig 13:**
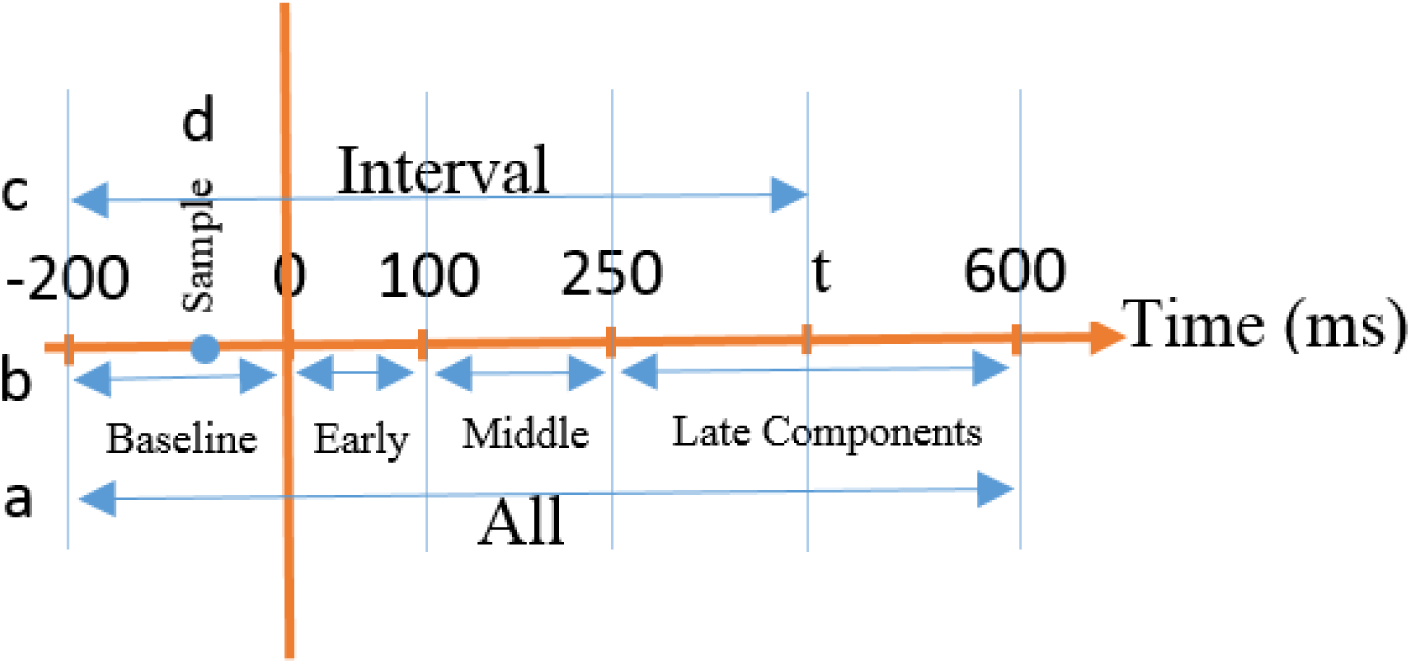
Different EEG/MEG temporal component selection regimes are used to define feature vectors as inputs to the decoder. **a)** All temporal components of a trial are used (All). **b)** Pre-defined temporal segments are used (Segments). **c)** Temporal components in the range of [‒ 200, *t*] are used (Intervals). **d)** Each single temporal point *t* is used (Samples).

At the end of each decoding analysis task, to judge the resulting R-squared values, we tested the hypothesis that *S_N ×p_* and *Y_N_* _× 1_ are independent of each other. This was done by making random permutations in the vector *Y_N_* _× 1_ and acquiring the R-squared value of the resulting regression each time as chance level. This analysis was performed for every subject and every task for each type of surprise.

### MEG temporal samples and segments

The definitions of the four regimes of temporal data selection are similar to those for the EEG analysis. The four basic time segments in each epoch of the MEG oddball task are defined as follows: from −200ms pre-stimulus to time 0 (Baseline), from time 0 to 100ms (Early Components), from 100ms to 250ms (Middle Components), and from 250ms to 600ms (Late Components).

## Acknowledgments

We thank Jennifer M. Walz and her colleagues for making the EEG oddball dataset publicly available [56]. We also thank Maxime Maheu and his colleagues for making the MEG oddball dataset and the Bayesian observer code publicly available [38].

